# Deep learning-based joint sequence–structure *de novo* membrane protein design

**DOI:** 10.1101/2025.08.15.670493

**Authors:** Lucas S.P. Rudden, Remo Bättig, Vinnie Andrews, Julie V. Nguyen, Martin Stoll, Lorenzo Scutteri, Michal Winnicki, Melissa J. Call, Matthew E. Call, Damien Thévenin, Patrick Barth

## Abstract

Deep learning has revolutionized soluble protein design, yet de novo transmembrane (TM) protein engineering remains hindered by scarce structural data, complex membrane-specific interactions and conformational dynamics. We developed TMDiffusion (TMDF), a joint all-heavy-atom sequence–structure diffusion model trained to capture the full interaction diversity of natural TM proteins, including weak and polar contact networks. TMDF designs diverse TM architectures—associating domains, inhibitors, and conformational switches—in a single step, achieving >70% experimental success. A crystal structure of designed proteins matches predictions with atomic accuracy. Leveraging TMDF, we built synthetic single-pass receptors whose de novo TM domains toggle between conformations, enabling precise control of signalling outputs consistent with predicted equilibria. These results show that membrane-adapted DL models can accurately encode and program TM association energetics and conformations. TMDF establishes a general framework for bottom-up design of TM proteins with programmable functions, advancing both mechanistic studies of membrane proteins and development of next-generation therapeutics.

## Introduction

Membrane proteins constitute ∼30% of the proteome and ∼70% of drug targets(*1*), mediating essential roles in cell communication, immunity, and neuronal signalling(*2–4*). Yet, they account for less than 1% of experimentally determined structures, reflecting persistent challenges in their structural characterization(*5*) and limiting advances in their prediction and design. Existing membrane protein design strategies have largely relied on knowledge-based approximations of the lipid environment(*6–10*) or on engineering membrane-compatible variants of hyperstable soluble scaffolds(*11, 12*). However, such approaches fail to capture the full chemical and structural diversity of transmembrane (TM) interactions and the delicate energetic balances that underpin their conformational dynamics(*13–15*). Unlike soluble proteins, TM domains are shaped by unique patterns of weak hydrogen bonds, polar networks, and solvent-mediated interactions—often within their hydrophobic interiors—that govern association equilibria and functional transitions. These interaction networks are enriched in backbone-side-chain atomic contacts (e.g. Gly-zipper motifs) that are very sensitive to the precise position of the backbone and amino-acid sequence, supporting the need for joint structure-sequence sampling. Current Deep Learning methods, such as RFdiffusion(*16*) and ProteinMPNN(*17*), sample these spaces sequentially, limiting their ability to model such interdependent features. As a result, the finely tuned energetics regulating receptor signaling and the activity cycles of channels and transporters remain largely inaccessible to current *de novo* membrane protein design methods.

*De novo* protein design holds transformative potential for drug design(*18–20*), biotechnology(*21, 22*), vaccine development(*23, 24*), and beyond. Artificial intelligence (AI) generative approaches, particularly denoising diffusion probabilistic models (DDPMs), have advanced both protein structure prediction(*25, 26*) and generation. Designing function, however, requires AI methods that jointly generate structure and sequence(*27*)—yet such approaches remain scarce. ProteinGenerator(*11*), for example, diffuses in sequence space while conditioning on RoseTTaFold2(*28*) structural outputs, but trails RFdiffusion in performance, likely due to the absence of explicit structure distribution and modeling(*29*). Other joint modeling frameworks— CarbonNovo(*30*), DiffAb(*31*), and MultiFlow(*32*) —omit side-chain atoms, neglecting critical biochemical determinants of folding and interactions. Protpardelle(*33*) performs Euclidean all-atom diffusion but predicts sequences using a separate network(*34*), providing transformers(*35*) in the denoising loop with only implicit sequence information. To our knowledge, no existing AI method explicitly represents and diffuses over the joint all-atom structure–sequence distribution, a capability that could capture the full complexity of protein energetics and enable conformationally dynamic protein structures and functional design.

## Results

### TMDiffusion, an all-atom sequence-structure AI design tool for TM proteins

To address these limitations, we developed TMDiffusion (TMDF), the first joint all-heavy atom sequence-structure AI design tool built specifically for TM proteins. To overcome limited training data, we restrict the learnable space to TM α-helical dimers—a motif common to all single- and multi-pass membrane proteins, including receptor tyrosine kinases and G-protein coupled receptors—allowing an impressive consideration of functionality to be integrated directly into the model (**Figure 1a**). Unlike most AI protein design tools, TMDF operates directly on a full all-atom description without tertiary neural networks or sequential post-processing. During forward diffusion, noised features are extracted distinctly from Euclidean (CA coordinates, amino acid identity, chain identity), SO(3) (backbone orientation), and SO(2) (sidechain dihedrals) distributions (see **Methods**, **Figure 1a**), while the novel denoising architecture is trained jointly on all inputs (**Figure 1b**). The learning of TM-specific biophysical interactions—including salt bridges, SmallxxxSmall motifs, CA–H···O hydrogen bonds, polar cores, and aromatic interactions—is reinforced via explicit contact maps extracted from the noised version of an input protein at time *t* that are then fed into the neural network alongside our protein description. As described below, this enables precise recapitulation of native-like TM interfaces and generation of novel, hyper-optimized motifs, such as multi-SmallxxxSmall configurations with enhanced CA–H···O bonding and complex polar networks unseen in natural proteins. TMDF addresses key challenges in membrane protein engineering, enabling the design of TM inhibitors, TM assemblies with diverse topologies, TM structures conditioned on arbitrarily mixed backbone scaffolds and sequence context and synthetic receptors with programmable function (**Figure 1c**). Additionally, the model supports protein switch design by propagating shared logits across successive diffusion steps. This coupling ensures that, at every stage of the denoising process, sequence sampling is conditioned on the simultaneous representation of multiple conformational states, rather than on post hoc consensus or averaging of final sequence probability distributions(*17*).

**Figure 1:**
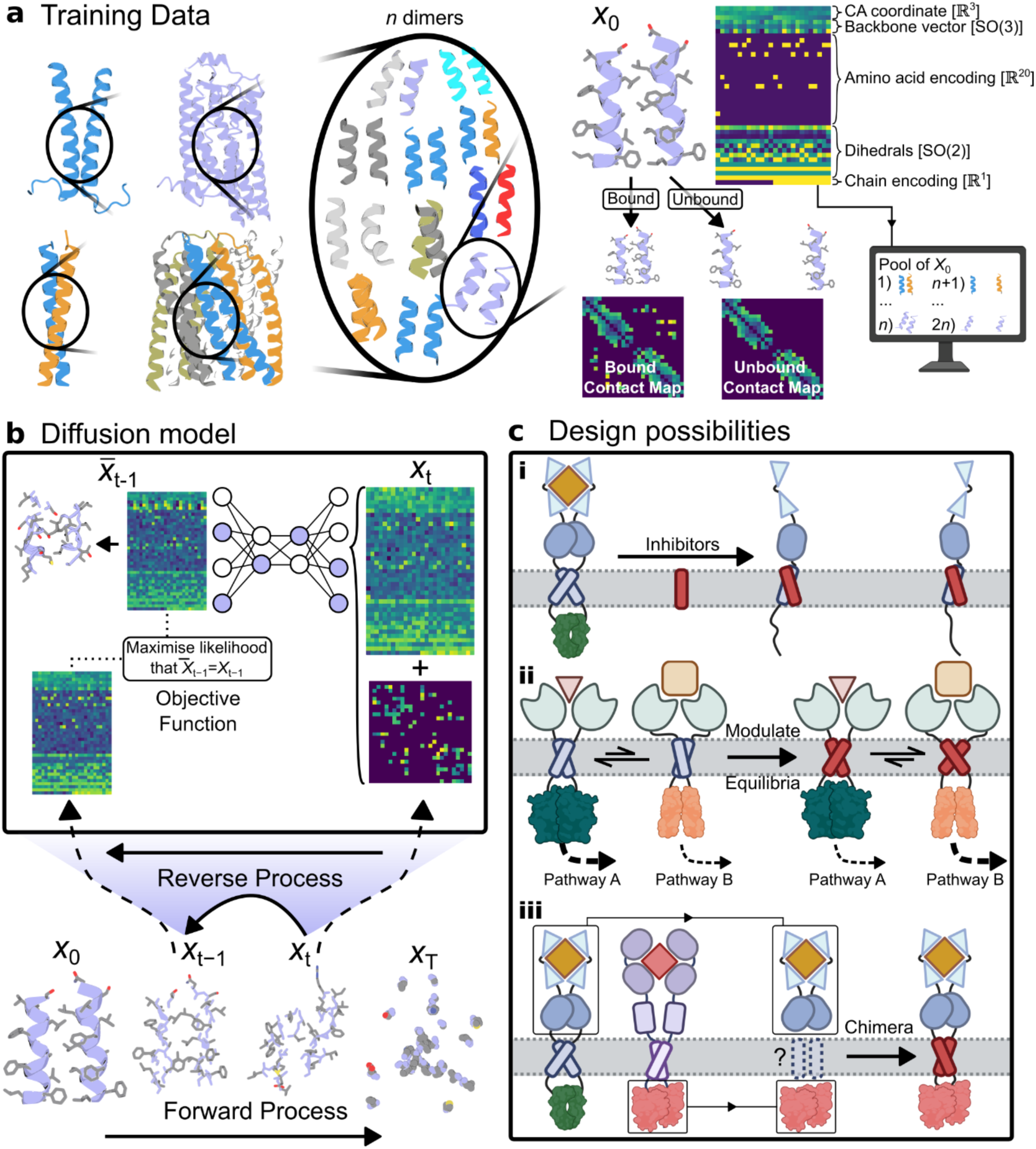
The TMDF model and applications. (**a**) Training data is made up of a large pool of homo/hetero dimers extracted from homodimers, heterodimers, multipass, and oligomeric TM proteins following a strict set of criteria (**Methods**). For each bound dimer/datapoint, we create a**n** unbound version to double the dataset size and generate a vectorised version of the protein as input for the neural network bearing the key features we want TMDF to learn. (**b**) During training, data is noised to time *t* and a contact map generated from the noised protein. These inputs are fed into the neural network, which predicts the *t* − 1 distribution of the input protein. The loss function maximises the likelihood that the neural network’s output and the true *x_t_* − 1 data match (**Methods**). (**c**) TMDF facilitates a large variety of design possibilities: (**i**) The design of inhibitors to outcompete native receptor association and function; (**ii**) The design of TM regions able to occupy two different conformational states, with modulated equilibrium to control the output signal/level of signalling; (**iii**) The construction of chimera that couple arbitrary EC and IC domains via *de novo* design of the TM domain. Schematic figures were created with BioRender.

### TMDiffusion recapitulates the sequence-structure distribution of TM α-helices

We first examined the joint sequence-structure space of TM α-helices learnt by TMDiffusion through the unconditional generation of 10000 designs and characterised their general properties. In summary we looked at the crossing angles **(Figure 2a**), general amino acid distribution (**Figure 2b**), the types of contacts formed by generated designs (**Figure 2c**), sequence/structure association (**Figure 2d**), and the dihedral angle distribution of raw outputs (**Figure 2e**).

**Figure 2:**
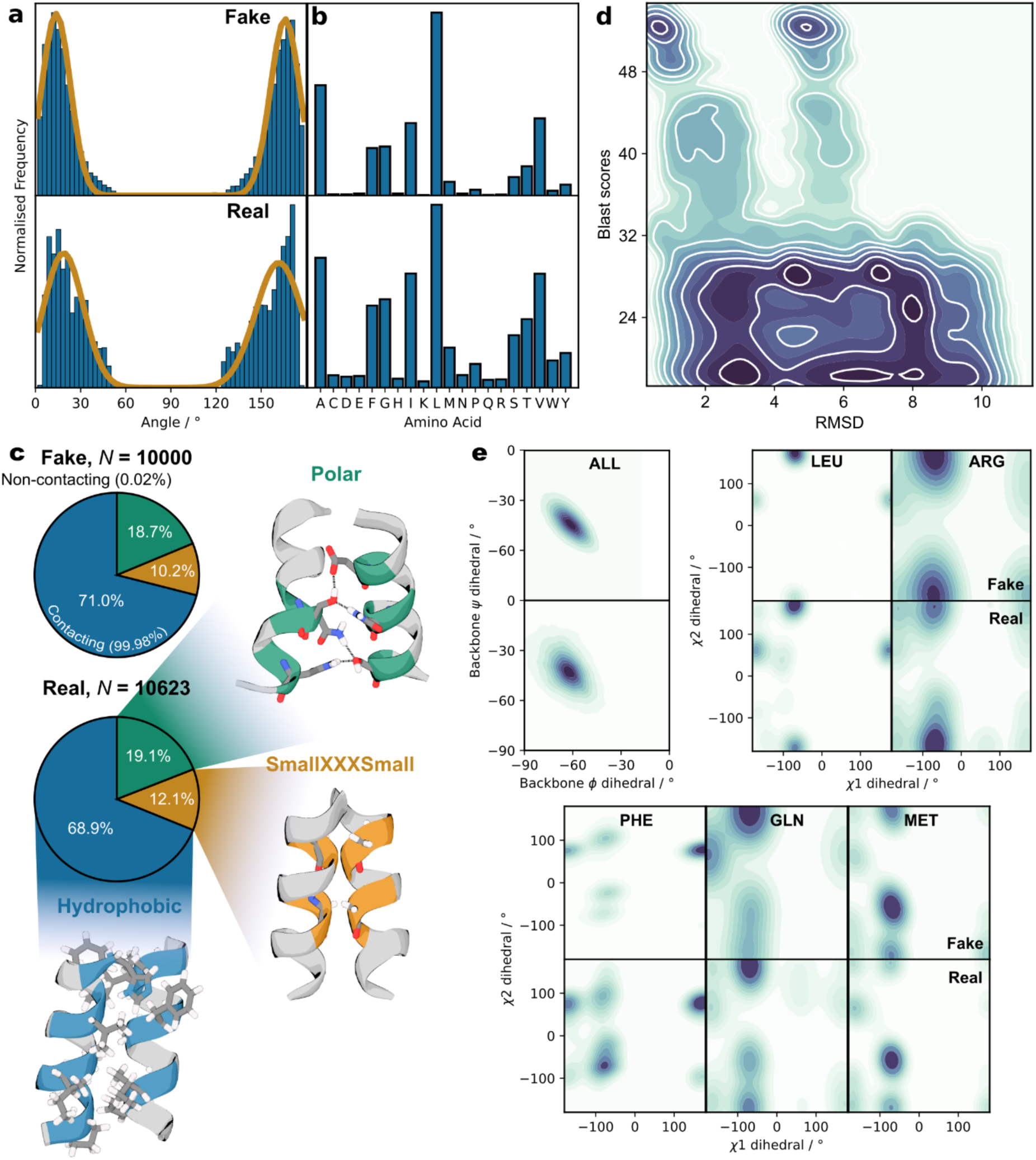
Learnt protein sequence-structure features. All key features that describe protein sequence/structure are learnt by TMDF, as shown by 10000 unconditional generated designs recapitulating the training data. (**a**) The crossing angle distribution of dimeric chains matches the real data distribution, albeit with some smoothing of the double Gaussian. (**b**) The amino acid distribution closely resembles the real data counterpart with some undersampling of polar amino acids. (**c**) Almost all generated designs bear a significant contacting interface, with motifs aligned with that seen in real TM proteins. (**d**) There is a close correlation between the RMSD and BLAST score between generated sample**s** and those in the training data. RMSD gives a measure of structural similarity, while BLAST corresponds to the sequence similarity. Their correspondence indicates that TMDF has learnt the sequence-structure relationship. (**e**) The Ramachandran space of both the backbone dihedrals, and the χ_1_ and χ_2_ sidechain dihedrals, closely match that seen in the training data.

The crossing angle influences the types of interactions between two helices, with different crossing angles favouring specific packing arrangements and sidechain interactions(*36*). Right-handed crossing angles, whether in parallel and antiparallel orientations, preferentially accommodate small residues spaced by three positions; forming the SmallxxxSmall motif, where the small residue is either glycine (predominantly), serine or alanine. Conversely, left-handed angles tend to favour small residues recurring every seventh position. Thus, the crossing angle serves as a key descriptor of the structural organisation of TM α-helical dimers. Figure 2a clearly demonstrates that TMDF accurately reproduces the general distribution of crossing angles observed in the training data.

The sequences of transmembrane proteins are fundamentally different to soluble proteins, with a large bias towards hydrophobic residues both in the protein core and surfaces facing lipids’ hydrocarbon tails. Despite a ∼5% underrepresentation of non-hydrophobic residues, TMDF recapitulates faithfully the amino-acid distributions in TM proteins (**Figure 2b**). While this bias could reflect overtraining, TMDF still incorporates non-hydrophobic residues strategically to optimise binding (see design case studies below), with a similar propensity of polar interactions and SmallxxxSmall motif insertion to the real data (**Figure 2c**). This indicates that their occurrence in TMDF’s output remains tightly coupled to their biochemical role. Indeed, when contact maps are omitted from the network, both the recapitulation of the amino acid distribution (**Figure S1b**), and the types of contact – be they hydrophobic, SmallxxxSmall or polar – are substantially diminished (**Figure S1c**). Thus, the apparent oversampling of hydrophobic residues may instead indicate that TMDF more readily identifies optimal subunit arrangements via hydrophobic packing, whereas the inclusion of non-hydrophobic interactions is more selective and deliberate, ultimately contributing to TMDF’s success.

Given the importance of learning the sequence-structure relationship of TM α-helices, we next assessed how sequences generated by TMDF relate to their predicted structures. For each generated sample, we compared its highest BLAST (sequence homology) score to any training sequence against the RMSD of the associated fake/real pair (i.e. a high BLAST score corresponds to high sequence similarity, **Figure 2d**). We observed a significant relationship between the two (i.e. sequences with high homology to the training data tend to also exhibit relatively low RMSDs), suggesting that the model has effectively learned sequence-structure relationship. In contrast, the ablation test without contact maps showed little correlation (**Figure S1d**), emphasising the role of residue-residue contacts in guiding learning. The density cluster at ∼5 Å RMSD coupled with high BLAST scores demonstrates that besides generating sequences folding into similar structures to that seen in the training data, TMDF can also generate putative alternative arrangements, an essential capability for designing conformational switches (see GHR case study below).

Finally, we explored TMDF’s ability to recapitulate the dihedral space of all amino acids (**Figure 2e**). The *φ* - *ψ* Ramachandran plot was faithfully recovered, an expected result given that all training examples occupy this narrow angle well. More notably, TMDF successfully regenerates the more intricate *χ*1 - *χ*2 landscape for nearly all amino acids, even when rarely observed within the probability distribution. Accurate rotamer placement is critical for capturing packing arrangements and thus the type of resulting biochemical interactions. While occasional steric clashes are observed in the output, the Ramachandran plots confirms that TMDF has clearly learnt the rotamer space, while Figures 2c/d show that it can then apply this knowledge correctly in a design context. Furthermore, the marginally worse Ramachandran plots observed in the contact map ablation case (**Figure S1e**), together with the results in Figure S1c/d, highlight the importance of the contact maps in enabling TMDF’s to learn the intimate relationship between nuanced biochemical interactions and the global backbone architecture required for precise packing.

### TMDiffusion generates high-affinity right-handed TM associations

#### We next sought to assess TMDF’s ability to build novel TM structures and binding

A large majority of TM dimer assemblies adopt right-handed topologies. A canonical example in this family is Glycophorin A (GpA), one of the tightest TM dimers ever characterized. GpA achieves high binding affinity through Gly-zipper motifs (GxxxG), providing 6 weak CΑ–H···O hydrogens bonds(*37*). Both the close packing arrangements, and these weak interactions, are critical to the stability of right-handed assemblies(*36*). We generated thousands of designs at different noising levels of the ground truth (PDB: 1AFO), from one or two single amino-acid substitutions to full destruction of the data, guiding the diffusion process via the GpA backbone, before filtering to a final selection of 16 designs based on a combined ColabFold(*38*)/Rosetta(*39*) scoring approach (**Methods**).

A diverse range of designs were produced featuring many novel and unusual interaction motifs. When guided by a ground truth, TMDF tends to produce solutions with higher sequence identity to the condition (**Figure 3a**), with an average sequence identity to the WT of 70%. However, most of the highest scoring designs when compared directly to WT sit within the low sequence identity tail of this distribution. Furthermore, while exhibiting similar global structures (< 3 Å backbone RMSD), designs are evenly distributed between WT-like and higher crossing angles (**Figure 3b**) with several optimal solutions lying beyond the naturally selected GpA WT features. Given that our selection criteria are agnostic to the input condition, TMDF achieves some optimised designs by bypassing high-energy physical barriers through one-shot generation, thereby identifying sequences far removed from the WT that are still able to fold into similar conformations.

**Figure 3:**
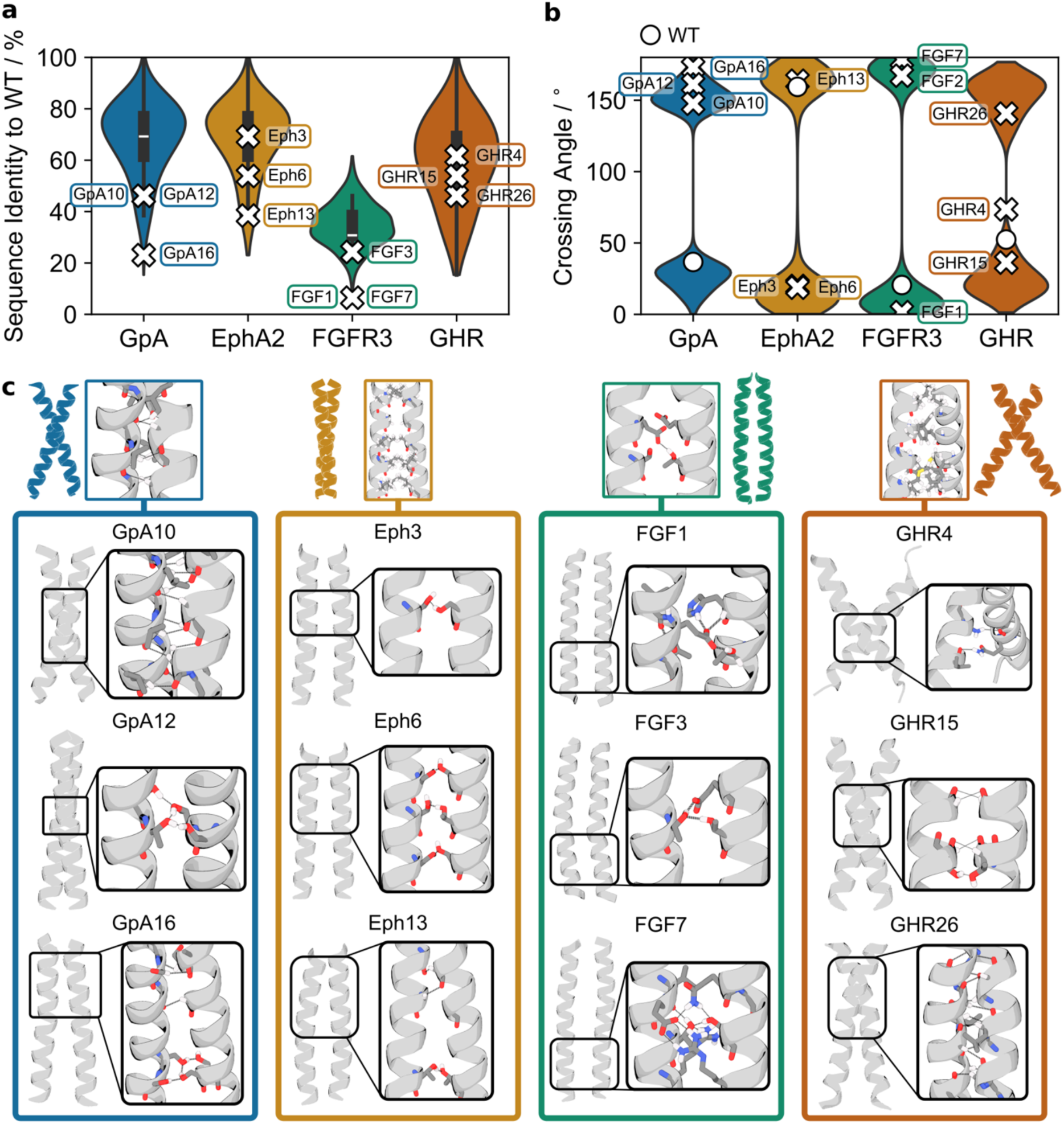
*De novo* design sequence-structure diversity. TMDF *de novo* designs novel and innovative motifs regardless of the design problem: GpA (right-handed helix), EphA2 (left-handed helix), FGFR3 (inhibitor design), GHR (two conformational states). (**a**) 10000 raw design output from TMDF. The sequence identity of generated designs to the WT depends on the design problem, with GpA/EphA2 featuring more similar sequences than the conformationally dynamic GHR, and FGFR3 where no condition was applied to inhibitor chain. (**b**) 10000 raw design output from TMDF. The crossing angle distribution sampled by TMDF again depends on the design problem, but in all cases a wide variety of angles are sampled by the neural network to achieve optimised binding. (**c**) A selection of unusual but often highly successful binding motifs sampled by TMDF for each design case – corresponding to the sequences mentioned in **a/b** and in Figure 4. Dotted lines correspond to H bonds, either between polar residues directly, or weak Cα–H···O bonds.

To determine whether these predictions hold true experimentally, we validated our selected designs using the *in vivo* TOXGREEN assay(*40*) (**Methods**), which quantitatively measures TM protein association in the inner membrane of *E. coli*. Immunoblotting confirmed that all designs, including the weakly dimerizing GpA G83I mutant, expressed and orientated correctly in the membrane. While GpA2 and GpA4 showed no significant binding, 3 designs performed similarly to WT (**Figure 4a**). The remaining 11 designs formed higher affinity oligomers than the WT, with GpA10, GpA6, and GpA12 yielding the highest signals, with GpA12 exceeding the WT by more than twofold. To validate the predicted binding interfaces of our strongly-associating TM oligomers, we next generated “poor” variants in which key designed binding interactions were disrupted by amino-acid substitutions. All GpA poor variants showed markedly reduced TOXGREEN signals compared with both the WT and the original designs (**Figure 4b**). For example, GpA10 dropped from 1.5x the WT signal to less than 0.5x, validating the structure of our designed binding interfaces.

**Figure 4:**
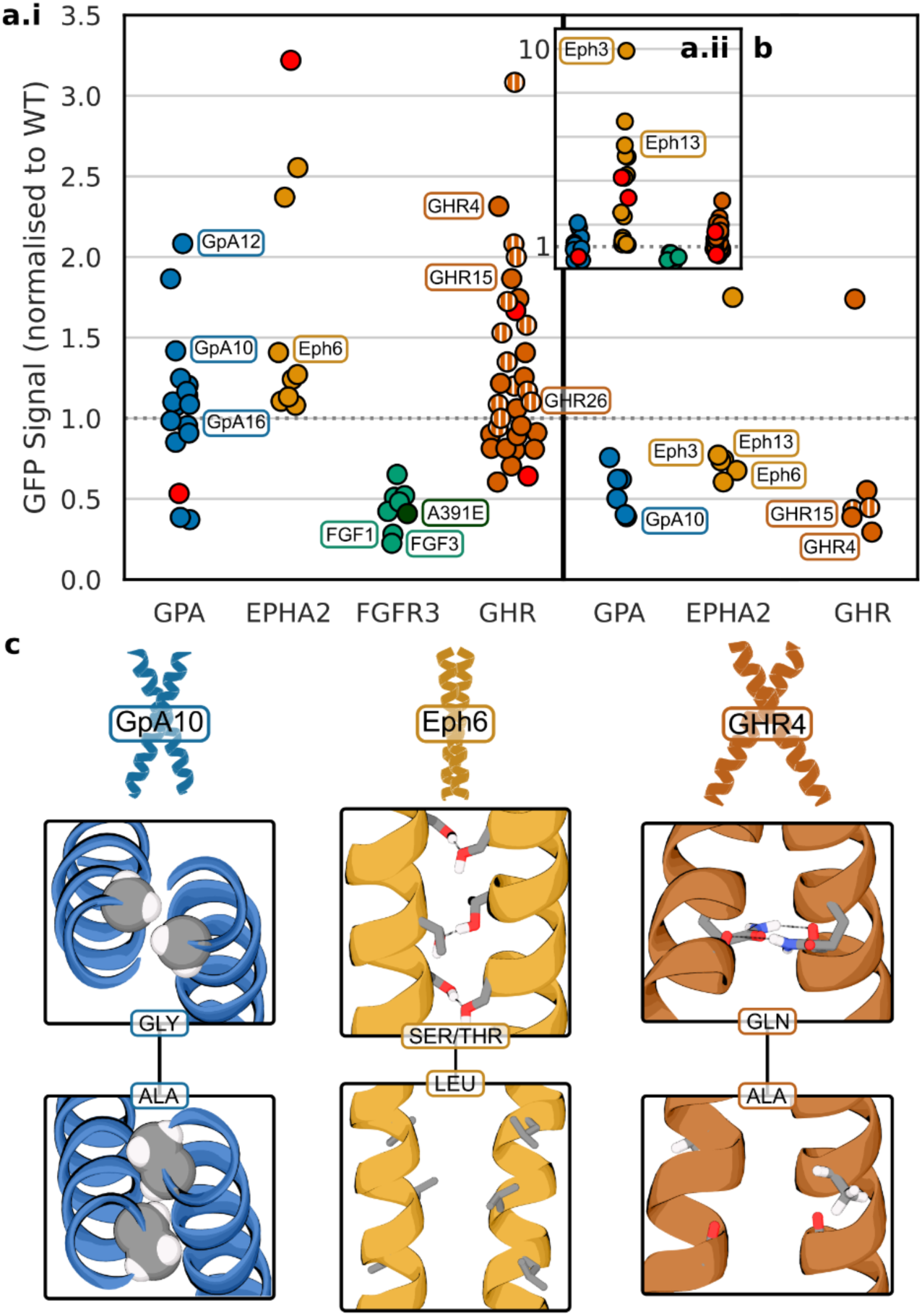
TM association propensity. TM oligomerization measured by TOXGREEN (DN-AraTM for FGFR3). (**a.i**) Average of TOXGREEN signals normalised against the WT (dotted line at 1). In FGFR3’s case, the DN-AraTM signals are normalised against the non-competitive ALA391GLU mutant versus empty, and the A391E mutant shown in a darker green. Red indicates supposed poorly oligomerising mutants extracted from the literature. With GHR, we indicate the first switch design round (GHR1-17) with filled circles, and the second design round where we optimized for the inactive state (GHR18-30) with striped circles. (**a.ii**) Inset showing a larger scale on the GFP signal axis to highlight the EphA2 designs significantly larger range of response. (**b**) TOXGREEN signals for the GpA, EphA2, and GHR sequences created to break key interactions at the designed interface. Only two out of the eighteen designs failed, indirectly validating TMDiffusionForge’s designs. (**c**) Visualisation of three purposefully damaged interfaces corresponding to the GpA10, Eph6, and GHR4 signals in **b**.

Exploring in more detail why some designs returned improved signals revealed a set of unusual interaction motifs, illustrating how TMDF can further optimise even highly stable dimers such as GpA (**Figure 3c**). **GpA10** hyperoptimises the close-packing interface by inserting a GxxxGxxxA motif, with the alanine positioned where the helices begin to diverge, better filling the inter-helical void. This design also programs **10 weak Cα–H···O hydrogen bonds**, a substantial increase over the WT’s 6. **GpA12**, in contrast, relies on an extensive polar network, with a chain of serine–threonine–threonine–serine hydrogen bonds providing strong intermolecular “glue.” While this arrangement leaves two unsatisfied polar groups on the serines, the stability conferred by these interactions is sufficient to drive the observed >WT oligomerisation. **GpA16**, although not outperforming WT, is highly divergent, sharing only 20% sequence identity. It combines a close-packing GxxxG motif (top of subfigure) with a polar network reminiscent of GpA12 (bottom of subfigure). However, its dimer asymmetry limits affinity—only three weak Cα–H···O bonds form—and the backbone curvature needed to accommodate both motifs likely imposes an energetic penalty, explaining its weaker TOXGREEN signal. Nonetheless, this unusual conformation demonstrates that TMDF can access diverse regions of the α-helical TM proteome to produce solutions that are distinct from the input sequence.

To further characterize the structure of the designed TM oligomers, six of the proteins were expressed in *E. coli* and purified. Five of six constructs exhibited the intended oligomerization state on SDS–PAGE gels (**Figure S4)** and all were reconstituted in lipid cubic phase for X-ray crystallographic analysis. Crystals of the GpA3 design diffracted to 1.7 Å resolution, yielding an atomic-resolution structure (**Table S1**). The X-ray model showed remarkable agreement with the TMDF prediction (Cα RMSD = 0.55 Å), with a large fraction of side-chain conformations predicted at atomic accuracy (**Figure 5)**. These findings validate TMDF’s capability to generate TM structures and interactions with atomic precision.

**Figure 5.**
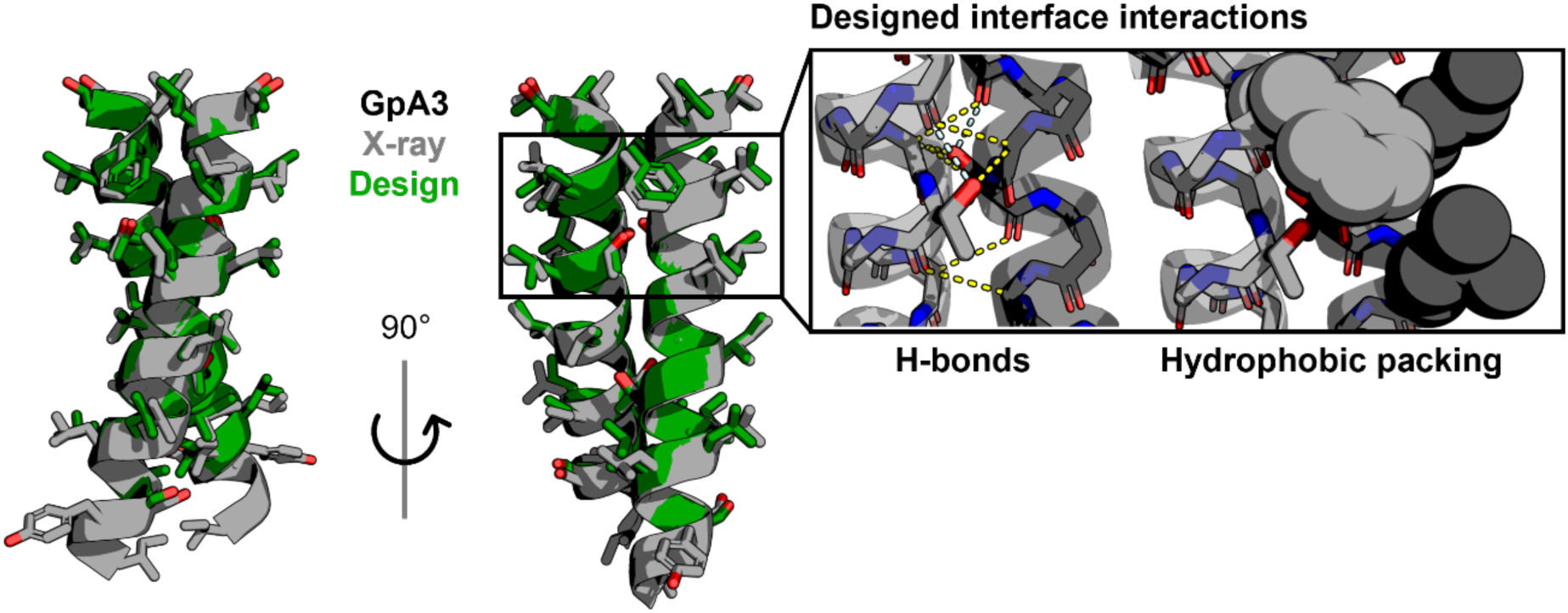
Structural characterization. Superposition of GpA3 designed model (green) and structure resolved by x-ray (grey) reveal sub-angstrom accuracy. The interface consists of polar, hydrophobic, as well as small (Gly) residues. The box highlights several of the designed interface interactions such as inter- and intrachain H-bonds (dashed lines), as well as hydrophobic packing (sphere representation).

### TMDiffusion can design left-handed TM oligomeric scaffolds

Parallel left-handed α-helical TM dimers are comparatively rare in nature and more difficult to model(*41*) due to their atypical topology and suboptimal packing arrangements. To test TMDF under these conditions, we generated left-handed designs, starting from that of Ephrin Type-A Receptor 2 (EphA2), whose binding is mediated by a heptad repeat of LxxxGxxAxxxVxxL(*42*) (**Figure 3c**). This required adjusting the design strategy– specifically, using AlphaFold2 for structure prediction instead of ColabFold which was unable to reproduce the WT fold, underscoring the complexity inherent to left-handed arrangements.

In contrast to GpA, high scoring designs were not biased towards lower sequence identities and alternative crossing angles (**Figures 3a/b**), and all final models featured a low RMSD to WT (< 1.5 Å). Expression and orientation were confirmed for all designs. Despite their close similarity to WT, improving binding proved markedly easier. Every design outperformed WT in the TOXGREEN assay, with one design – Eph3, featuring a 10x improvement over WT (**Figure 4a.ii**). Indirect validation of the dimers using disruptive mutations demonstrated that only one design, Eph11, did not adopt the expected conformation (**Figure 4b**).

In Eph6’s case for example, we disrupted the dimerisation interface through the swapping of the serine/threonines to leucine. These polar residues are clearly key to the dimer stability, as their replacement reduced the signal to almost half that of the WT – itself a weak oligomer. Overall, these results demonstrate that TMDF can design dimers within the more challenging left-handed α-helical TM space, and indeed easily improve on native TM interfaces.

A closer examination of the top designs reveals a prevalence of polar interactions inserted at the interface (**Figure 3c**). Unlike the multi-residue interaction networks typical of right-handed scaffolds, these consist of single serine–serine or threonine–threonine pairs. The best-performing design, Eph3, contains only one such insertion, consistent with the strong effect of optimally positioned polar contacts at left-handed TM interfaces(*43, 44*). Eph13 is unusual in that it maintains a left-handed orientation while introducing an asymmetric GxxxG motif, enabling two weak CA–H···OH hydrogen bonds—a motif not normally observed in left-handed dimers. In addition to a polar interaction near the C-termini, this atypical arrangement yields a GFP signal nearly sixfold higher than WT, illustrating how TMDF can repurpose interaction motifs from diverse structural contexts and deploy them in novel ways to optimize oligomerization.

### TMDiffusion designs potent TM inhibitors

Beyond designing optimized dimers from native scaffolds, we also sought to explore the therapeutic potential of TMDF. Several receptor missense variants with point mutations in the transmembrane region are linked to severe diseases(*45, 46*), often by disrupting the conformational equilibrium or allosteric transitions that regulate activation. One well-known example is the Ala391Glu mutation in the transmembrane domain of fibroblast growth factor receptor 3 (FGFR3), which causes Crouzon syndrome(*47*) —a genetic disorder characterized by craniosynostosis. The A391E mutation leads to receptor overactivation, and modulating its signaling has been proposed as a strategy to alleviate neurological complications associated with the disease(*48*). Biophysical studies indicate that this mutation enhances FGFR3 dimer stabilization(*49*). Motivated by this mechanism, we initiated a TMDF-based design campaign to generate competitive inhibitors targeting the mutant FGFR3.

Given an MD-relaxed structure of A391E FGFR3 (**Methods**), we tasked TMDF with freely diffusing an inhibitor TM chain against the mutant monomer. The network produced a set of designs with substantially lower sequence identity to WT than observed in the GpA and EphA2 design campaigns (**Figure 3a**). While this is not unsurprising given no bias was applied towards the mutant sequence for the designed chain, a measurable sequence homology to FGFR3 remained, underscoring the strength of TMDF’s learnt-sequence structure relationship. Despite the freedom to design the second chain without any bias, the crossing angle distribution of the designs was narrower than for GpA and EphA2 (**Figure 3b**). This restricted geometry likely reflects the packing challenge posed by the glutamate substitution, for which only a limited set of packing arrangements can effectively accommodate the anionic side chain. These results indicate that TMDF incorporates an implicit awareness of the lipid environment when making design choices.

A variety of complex interface arrangements were generated to satisfy the glutamate, ranging from single to multiple polar residues. While designs incorporating multiple polar groups often lacked sufficient hydrophobicity to fold within lipid membranes, seven designs expressed successfully and adopted the correct membrane orientation. We assessed heteromeric binding to FGFR3 A391E using DN-AraTM, a dominant-negative version of TOXGREEN. In this assay, hetero oligomer formation reduces GFP signal. All designs showed significant interaction with FGFR3 A391E and, except for FGF11, matched or exceeded the binding strength of the target’s strong self-association. Two designs could qualify as inhibitors by outcompeting the target self-association. For example, FGF3 used a single serine to engage the glutamate, whereas FGF1 employed both a histidine and threonine to satisfy the carboxyl group (**Figure 3c**). Overall, TMDF achieved a remarkable 47% success rate, despite numerous polar interactions at the TM interface, highlighting its potential for addressing challenging inhibitor design problems.

### TMDiffusion enables programmable control of single-pass receptor signaling

Finally, we sought to challenge TMDF with one of the most difficult problems in TM protein engineering: designing single-pass signaling receptors. In this receptor class, the relationship between conformation, binding, and activation remains poorly understood. Proposed activation mechanisms range from monomer-to-dimer transitions, to conformational switching between inactive and active states within preformed dimers(*13, 50*). For example, NMR studies of the Growth Hormone Receptor (GHR) TM domain have identified two distinct dimer conformations—attributed to inactive and active states(*51*)—whereas studies of the full-length receptor support an activation mechanism driven by a monomer–dimer equilibrium(*52*) (**Figure 6a**). To test these possibilities, we tasked TMDF with designing sequences that either favored one of the dimer conformations or switched between them (**Figure 6b**). We reasoned that if GHR activation occurs through conformational changes within preformed dimers, signaling should be sensitive to the conformational switching properties of the TM domains. Conversely, if activation follows a monomer-to-dimer mechanism, signaling should depend primarily on dimerization propensity rather than on the ability to switch conformations.

**Figure 6.**
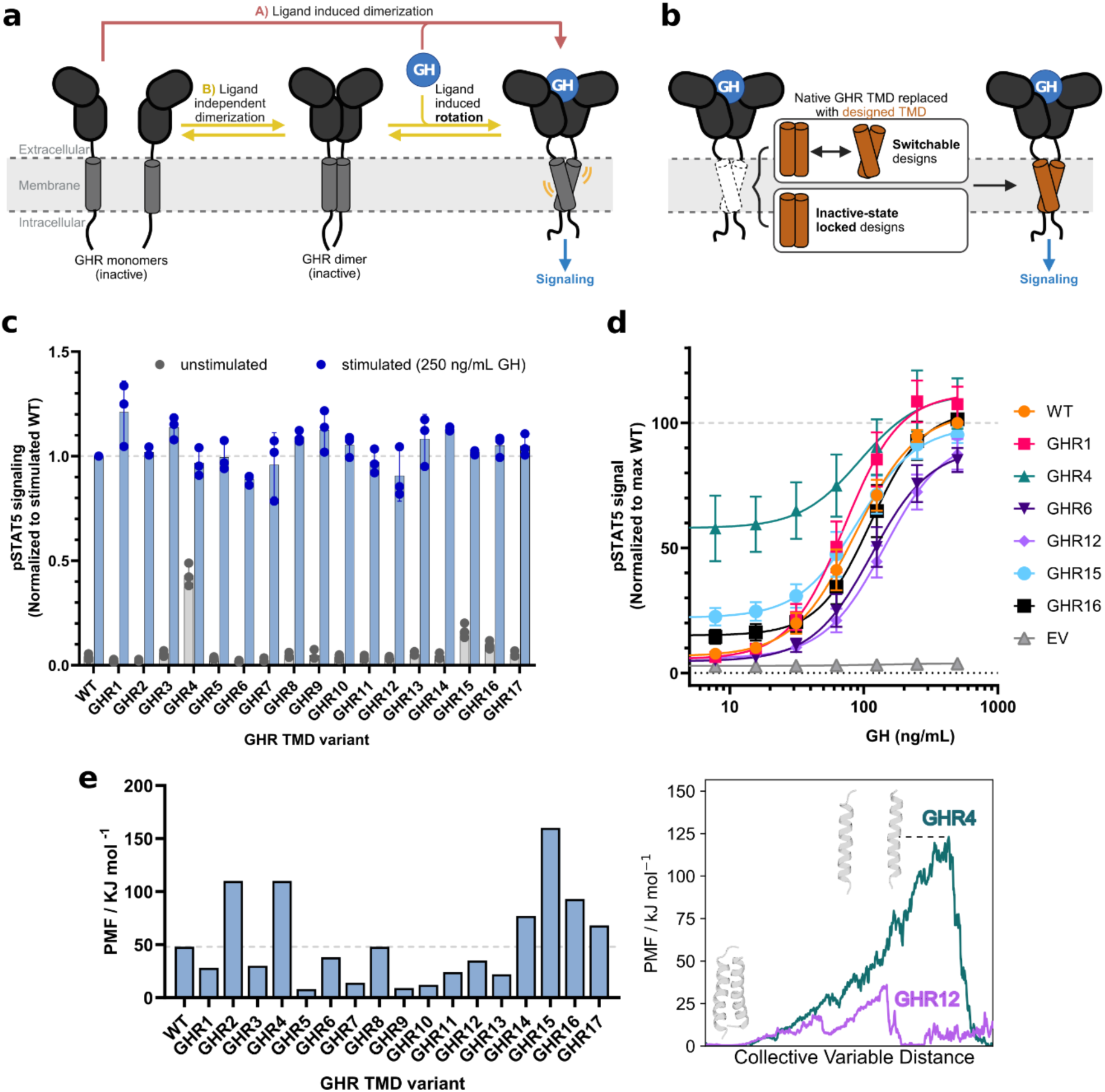
Synthetic GHR receptor signaling. (**a**) Schematic representation of two putative mechanisms of growth hormone receptor activation. A) Ligand binding induces dimerization, resulting in receptor activation. This mechanism lacks an inactive dimer state. B) Two monomers can undergo ligand-independent dimerization, resulting in an inactive dimer. Upon ligand binding, conformational changes result in switching of the TMD interface and receptor activation. (**b**) To probe GHR activation mechanism, the native TMD of full-length human GHR was replaced by designed switchable or inactive-state locked TMDs. Constructs were experimentally characterized in HEK293T cells to assess ligand-induced pSTAT5 signaling (panels **c/d**). (**c**) Comparison of basal activity and ligand-induced pSTAT5 signal of GHR variants, *n* = 3 biological replicates. (**d**) Dose response curves, of selected GHR variants, *n* = 3 biological replicates. (**e**) (**Left**) Maximum energy barriers observed for either active or inactive state corresponding to **Table S2**. Higher potential of mean force (PMF) tends to correlate with higher measured basal activity. (**Right**) Example of steered MD PMF profiles for designs GHR4 and GHR12. The barrier denotes the point at which the two monomeric units separate, its inverse corresponds to the free energy of binding for the dimer. Schematic figures were created with BioRender.

For this task, we challenged TMDF to determine whether it could account for multiple conformational states simultaneously during design and directly optimize one state over the other. Unlike existing AI-based multi-state design strategies, which impose switching only after sequence generation by identifying overlaps in the final sequence pool, our method integrates sequence and structural information for both target states at every diffusion step, embedding state-selective optimization into the design process itself (**Methods**).

In this switchable mode, the generated sequences were substantially more diverse relative to the WT than in the comparative GpA or EphA2 design cases (**Figure 3a**), despite conditioning on the same sequence in both states, and they exhibited a broader crossing-angle distribution overall (**Figure 3b**). This indicates that TMDF adopts a more exploratory strategy, sampling further from the WT to accommodate both states.

We selected 17 sequences predicted to fold into both inactive and active states (solid circles in **Figure 3a**), and 13 sequences optimized for the inactive conformation (striped circles in **Figure 3a**), for experimental characterization. All designs were expressed and oligomerized in the cell membrane, with 19 outperforming WT GHR (**Methods**). Among these, the switchable designs GHR4 and 15 showed both strong association and preference for respectively the inactive or active form as assessed by steered MD simulations (**Figure 3c**, **Figure 6e**, **Table S2**). The relative magnitude of TOXGREEN signals between GHR1-17 and GHR18-30 (**Figure 3a**) also indicates that when just one state is optimised with TMDF, resultant designs have a significantly higher oligomerization propensity, thus suggesting that targeting multiple states during design may require a compromise with binding affinity. Mutating out the key binding residues in several designs validated their predicted binding interface and our steered MD analysis, with markedly reduced TOXGREEN signals (**Figure 3b**).

We next examined how the equilibrium and conformational switching properties of these designs influence GHR signaling. The native TM region was replaced with the *de novo*– designed domains, and signaling responses were measured in HEK 293T cells. All engineered receptors responded robustly to GH, with ligand-induced activities ranging from 86% to 120% of WT but exhibited notable differences in basal activity (0.5x – 10x of WT) (**Figure 6c/d**). These differences did not correlate with conformational switchability or state preference; however, basal activity displayed some correlation with TM domain dimerization propensity, suggesting that the monomer–dimer equilibrium may act as a potential driver of receptor signaling. Together, these results support a model in which GHR activation is primarily driven by oligomerization, with the active state sampling a broad conformational space compatible with multiple TM interaction surfaces. They further highlight the broad utility of TMDF for building single-pass receptors with finely tuned, programmable structures and signaling properties.

## Discussion

Deep learning approaches have made great strides in soluble protein design but have not been explicitly developed to address the unique challenges of membrane protein design— namely, the scarcity of experimental datasets, high conformational flexibility, and the delicate balance of molecular interactions governing folding and assembly. To overcome these hurdles, we developed TMDF, a joint all-heavy-atom sequence–structure diffusion model trained on transmembrane (TM) helical dimers extracted from membrane protein structures.

TMDF achieved unprecedented recovery of sequence–structure motifs enriched at TM interfaces, demonstrated a high success rate in designing TM structures, inhibitors, and conformational switches, and enabled the creation of single-pass receptors with programmable control over signaling functions. In its current form, TMDF offers broad utility for diverse challenges, including the design of single-pass biosensor receptors for therapeutic cell engineering, precision TM mini-antibody therapeutics, and the mechanistic exploration of membrane protein functions.

Despite its strong performance, the approach could be further improved by enhancing stereochemical accuracy and protein length control. Recurrent steric clashes in generated structures stem from modeling contacts via residue centers of mass. Beyond TM dimers, assembling multiple, partially overlapping dimer interfaces could enable programmable control over local stability and switchability, key features for complex multi-pass membrane protein design.

Overall, our study demonstrates that Deep Learning models specifically adapted to membrane proteins can accurately recapitulate and program TM structures and associations. TMDF establishes a general framework for the bottom-up design of TM proteins with programmable functions, expanding the reach *de novo* protein design using Deep Learning into the largely unexplored membrane protein universe.

## Methods

### Computational Methods

#### Training data

The 190 PDB homodimeric TM dimers structurally available at the time of dataset acquisition (September 2021) were inadequate to train any diffusion model for our design purposes. Expanding beyond this dataset, the 1038 heterodimers were also insufficient. We, therefore, leveraged the much greater reservoir of 5379 multipass membrane proteins in conjunction with the existing homo- and heterodimeric datasets to curate our training data. Specifically, we extracted dimeric binding motifs from the multipass membrane proteins analogous to our existing dimers, in other words not involved in tertiary interactions with neighbouring regions. Our rules for selecting training samples from the multipass set are as follows:

1. Overall resolution must be < 3 Å. This is true also for the homo- and heterodimers.
2. All residues must be buried within the membrane, as determined via the Orientations of Proteins in Membranes database(*53*).
3. Any gaps less than 2 residues were repaired using Modeller(*54, 55*); otherwise, the sample is dismissed.
4. Both subunits must be *α*-helical (determined via DSSP(*56*)), with a permitted buffer of 2 residues on either side of the helical motif.
5. A helix moving through the membrane must contain at least one residue in contact with another from ***one*** other helix within a cutoff of 5 Å (ignoring hydrogens)(*57*). With the contact site(s) defined as the midpoint of the helix-helix interaction, we move stepwise up and down from this midpoint until 13 residues have been selected in total for the interacting motif. This approach is the same for both chains of the ‘dimer’.
6. Based on the 5 Å cutoff, the sample is dismissed if any residue in either helix is in contact with a tertiary subunit.

All biomolecular manipulation was performed using the biobox(*58*) Python package. In the case of redundant structures deposited under different accession codes, we take only the highest resolution case. With NMR, we extract only one conformation of the ensemble. These measures ensure that we avoid overfitting during training.

Our final database consists of 10623 individual dimeric fragments. We then augment this to 42492 total samples by swapping the order of chains A and B and creating a labelled dataset consisting of the fragments in a bound and unbound state. This entire pipeline results in a split of 33.4% and 66.6% parallel and anti-parallel samples, respectively. Finally, we use Rosetta to idealize and relax our fragments using the high-resolution membrane scoring function to ensure our residues are well-packed. The backbone is kept frozen throughout the simulated annealing.

### Protein mapping

The 26 total residues of each dimer are converted into a one-dimensional tensor that is effectively a one-to-one mapping of the protein. We treat each residue as its own canonical orientation frame based on a plane defined by the backbone N, CA and C atoms. The remaining backbone and sidechain atoms can then be derived from these frames using both the torsion angles of the protein and a one-hot encoding of the amino acid identity. We also include a chain-based positional encoding such that the model recognises the respective chains A and B, and in theory, *M* chains.

In summary, given a 26-residue protein, we learn a generative prior over the following features:

1. 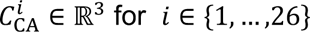, the Cartesian coordinates of the CA backbone atoms.
2. *V*^i^ ∈ SO(3), the unit vector defining the global rotation of the N, CA, C centred at 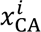.
3. *A^i^* ∈ {1,…, 20}, the categorical distribution of amino acid identity of the *i^th^* residue.
4. 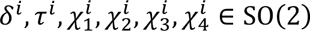, where *δ* is the dihedral through N, CA, C, O; *τ* the dihedral O, C, CA, CB; and the four *χ* angles that define the rotamer state connected to the *i^th^* CA. We drop χ_5_ as it only applies to arginine which infrequently appears in membrane proteins, and in the small number of cases it does appear, predominantly equates to an angle of *π*. We treat these dihedrals, *θ*, as the Pythagorean trigonometric identity along the feature vector.
5. *r^i^* ∈ {1,2}, the chain identity of the *i^th^* residue.
6. 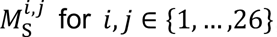, the sidechain – sidechain contact map, where we take the centre of mass of the sidechain (CA in glycine) as out contact point.

The contact maps are not required to regenerate the protein since a full description is provided by the preceding five features. Instead, they are used purely to emphasise residue contacts in the transformer.

After centering all fragments to the origin via their centres of mass, we normalise all Cartesian coordinates of the dataset between −1 and 1 based on the most extreme coordinates, retaining the normalisation constants for generation time.

We perform backmapping to the 3D structure of the protein in an entirely differential manner such that relevant biophysical quantities calculated during training (see neural network summary below), can be backpropagated against. We first obtain the CA coordinates of each residue simply based on the unnormalised values of 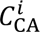 and apply the respective chain identities. We next regenerate the N, CA, C canonical frame from the global rotation using the reverse Gram-Schmidt process. Specifically, within the context of our canonical frame, given a rotation matrix *R^i^* with columns 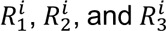, and translation *T^i^* that represents the position of the 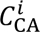 coordinates:

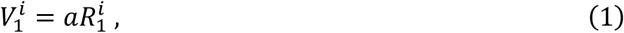

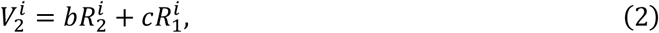

where *a* = 1.523 Å, *b* = 1.36 Å, and *c* = 0.52711 Å, are all constants based on the internal bond lengths within the canonical frame. From this, we find our coordinates for the N–CA–C triangle of residue *i*:

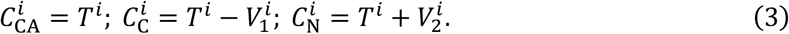

The forward Gram-Schmidt process is used when vectorising the proteins for the training data:

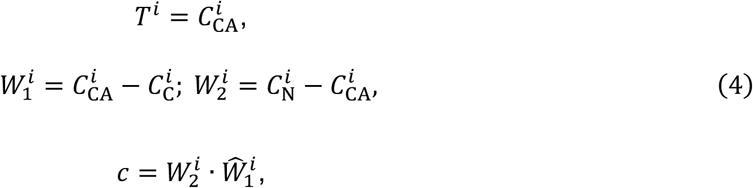

where |·| denotes the dot product, 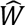 the unit vector, and *c* is the same constant as above.

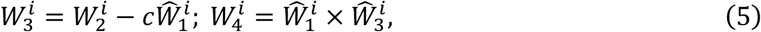

where |×| represents the cross product. Finally, we obtain the 3 × 3 rotation matrix:

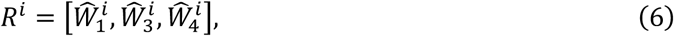

where the vectors represent the columns.

When extracting the sequence, we apply the Gumbel softmax trick(*59*) to our 1-hot encodings, which at time *t* of the diffusion process represents a series of probabilities for each amino acid. This pseudo-quantum mechanical property within our diffusion model means we retain a superposition of sequence identity states that collapses once measurement (converting back into a 3D structure) occurs. Therefore, this collapsed sequence ‘wavefunction’ represents the most probable singular sequence and structure identity at any diffusion timepoint. Thus, a powerful consequence of this fact is that the model can adapt its design choices on the fly based on optimising the sequence-structure combination during generation, effectively providing full flexibility to the protein backbone while designing the sequence.

Following the collapsing of the sequence identity, we attach the relevant sidechain to each residue from a library, extract the dihedrals and apply them to the stitched residues.

### Diffusion model

Denoising diffusion probabilistic models (DDPM)(*60*) are a class of generative models that aim to model the data generation process as a discrete iterative denoising process from a random prior. The process of moving from this random prior to a clean, *de novo* sample is the backwards process. Training our model begins with the forward diffusion process, wherein a drawn clean sample, *x*_0_∼*p*, drawn from our distribution of dimeric fragments *p*, is corrupted via a noise process defined in terms of a stochastic differential equation (SDE) of the form:

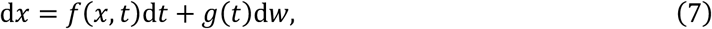

where *w* is a Brownian process on our manifold (either Euclidean, SO(3) or SO(2)), *f*(*x*, *t*) is a drift term, and *g*(*t*) a diffusion term at time *t*. In applying the noise to the forward process, we elected to use a variance exploding SDE as it: 1) enabled greater variance in the various mean posterior for the different manifolds we were operating over (where it is necessary for SO(3) and SO(2)), and because of these different manifolds, 2) facilitated simpler formulation of noise scheduler synchronicity between the different distributions, particularly SO(3) and SO(2) which are uniform.

The variance exploding SDE is defined as:

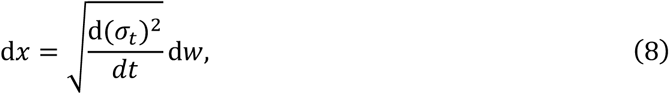

where *σ_t_* is a hyperparameter representing the amount of noise to be added according to the ***noise scheduler***, and d*w*∼*N*(0,1) provides the random noise to be added at timestep *t*. Given the approximation that the continuous diffusion process can be described through a discretised process over a large but finite number of steps, and using the Euler-Maruyama equations(*61*), we can write our discretised SDE as:

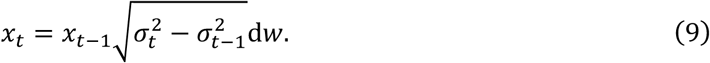

The 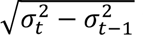 term represents the transition kernel since:

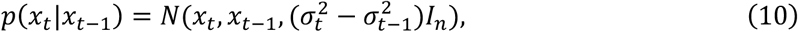

where *σ_t_* is the marginal:

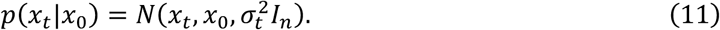

Moving forward, we will represent 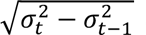 as *ε_t_*, and refer to it as the noise scheduler. Equation 8 is described as a variance exploding SDE owing to sample *x_t_* being extracted from an increasingly large multidimensional Gaussian.

Based on this suitable form of the SDE, the forward diffusion process can be treated as a Markov process:

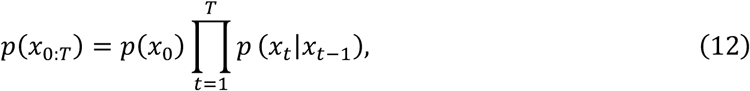

with transition kernel *p*(*x_t_*|*x_t_*_-1_) equating to Equation 10.

The core concept behind DDPMs, is that we can introduce a reverse Markov process defined via the variational transition kernel *p*(*x_t_*_-1_ |*x_t_*):

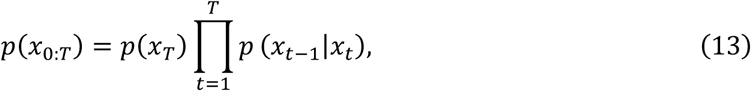

where any distribution appropriate to our manifold could be selected to parameterise this inverse transition kernel. For reasons discussed below, for our Euclidean data we adopt a normal distribution, for our 3D vector data an Isotropic Gaussian on SO(3), and for our dihedrals a Wrapped Normal distribution on SO(2). In the case of our *Dφ* distributions for SO(3) and SO(2) (we will use *φ* to describe both), our inverse transition kernel becomes:

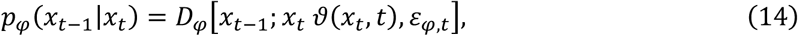

while for the Euclidean distribution it becomes:

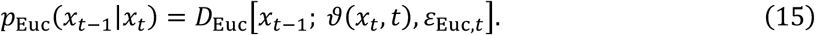

*υ* is a neural network predicting the residual or dihedral rotation to apply to *x_t_*, or predict *x_t-_*_1_ directly in the Euclidean case, and thus provides the mean of the reverse kernel. *ε_i_*_,*t*_ is the appropriate noise scheduler which depends on the manifold, where *i* represents either *φ* or Euc, which we typically simplify to just *ε_t_*. Note that *θ* is a ***shared*** neural network between all data types (hence the lack of subscript) that receives all *x_t_* and all *ε_t_* simultaneously, even if they are treated individually during the forward diffusion process. *θ* also predicts the variance of the reverse kernel via *ε_t_*. The output of the SO(3) and SO(2) data from *θ* are parameterised in their respective Lie algebra forms so(3) and SO(2), which we also use for input into the neural network.

Provided this reverse Markov process can be successfully trained to match the forward process, novel samples can be generated from *p*_0_ by initialising from the random prior at *p_r_* and iteratively sampling from the reverse kernel *p*(*x_t_*_-1_ |*x_t_*).

In DDPMs, the typical approach to train the individual transitional kernels is by approximating the reverse Markov process by minimising the evidence lower bound. Ho *et al.*(*60*) demonstrated that the variance over this loss could be reduced by using a closed form expression of the reverse kernel *p*(*x_t_*_-1_ |*x_t_*, *x*_0_) conditioned on *x*_0_, enabling the evidence lower bound to be rewritten as Kullback-Leibler divergences between Gaussian transition kernels. In other words, they were able to optimally learn the denoising process by predicting the noise added in the forward process and minimising the average squared distance between the predicted *x_T_* and the ground truth at each timepoint. Our distributions on SO(3) and SO(2) provide no equivalent closed form expression of the reverse kernel that Euclidean DDPMs do. Therefore, based on work by Jagvarel *et al.*(*62*), we consider an alternative expression for the evidence lower bound:

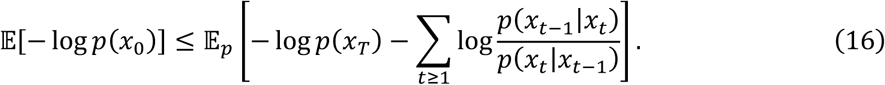

We optimise this by maximising the log likelihood of individual transition kernels log *p*(*x_t_*_-1_ |*x_t_*) from a noised state at time *t* to a denoised state at *t* − 1. In other words, during each iteration of training we simulate the forward Markov process before asking the network to predict *t* − 1, and calculating the log likelihood of that prediction belonging to the true *t* − 1 distribution using the following loss:

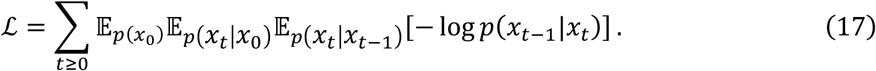

In a practical sense, we sum over the individual contributions from our protein mapping features (see above) to provide the loss.

### Data distributions

Owing to the different manifolds our data exists on, we must adopt three alternative distributions, *D*, to sample from in Equations 14 and 15. Our Cartesian CA coordinates, amino acid identities and chain identities exist in Euclidean space, on which the solution to the diffusion process is a Gaussian(*60*), parameterised by a mean, *μ*, in our case either the Euclidean coordinate or identity encoding, and scalar variance *ε*:

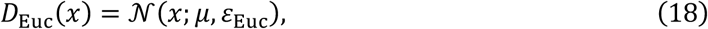

where *x* ∈ ℝ*^n^*. *ε*_Euc_ is the general form of the noise scheduler which we mentioned above and discuss in more detail below.

In contrast, for the compact manifold on which SO(3) exists the solution takes the form of the Isotropic Gaussian distribution(*63*), ℐ*G*_SO(3)_(*γ*, *ε*_SO(3)_), an infinite series(*64*) parameterised by a mean rotation, *γ*, and a scalar variance *ε*_Euc_ similar, but not identical, to *ε*_Euc_ for the Euclidean manifold:

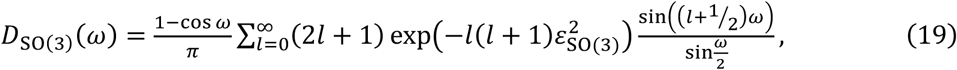

where we write the distribution in an axis-angle form, with *ω* ∈ [0, *π*] and the term 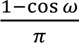 used as a scaling factor. In a practical sense within our diffusion model, *ε*_SO(3)_ is always smaller than 1. Matthies *et al.*(*65*) demonstrated that a closed-form expression can approximate Equation 19 when *ε*_SO(3)_ < 1:

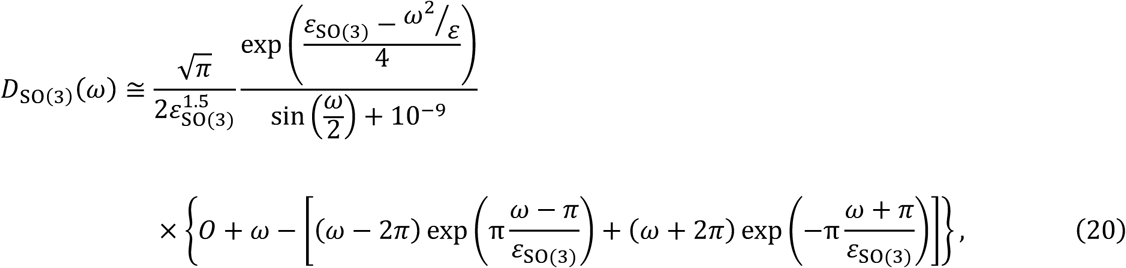

where *O* is defined as:

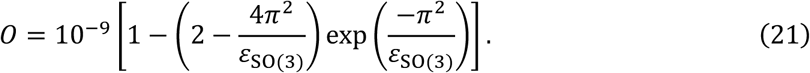

We include the *O* and 10^-9^ terms to avoid issues with numerical precision during back propagation.

Finally, we sample our SO(2) dihedrals using the Wrapped Normal distribution as provided by Mardia(*66*). Similar to Equation 19, the initial probability density function takes the form of an infinite series owing to its wrapped nature:

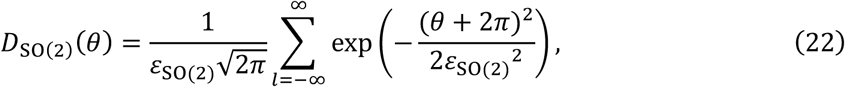

where *θ* ∈ [0,2*π*]. Based on a fundamental property of wrapped distributions shown by Mardia, namely that the Fourier transform of the general form of Equation 22 is integrable, we can obtain a more useful representation:

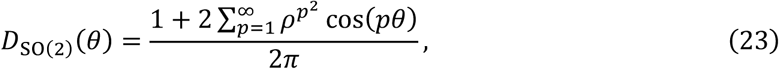

where *ρ* is defined as:

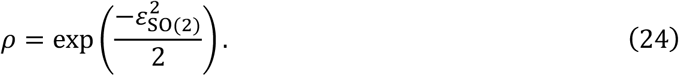

As *ρ* → 0, *D*_SO(2)_(*θ*) tends to a uniform distribution, while as *ρ* → 1 it focuses to a Dirac delta. For practical purposes, the first three terms of Equation 23 suffice. We elected to use the Wrapped Normal distribution and not the Von Mises, Wrapped Cauchy, or Cardioid distributions owing to its full width at half maximum striking a balance between sufficient sampling away from the mean without approaching the uniform distribution too quickly.

### Noise scheduler

The noise scheduler, *ε*, dictates the amount of noise added during training and, fundamentally, describes the variance of our distribution at any time point. Specifically, at training time the noise scheduler defines the variance of the distribution at each timepoint, from which we then extract our noised data at time *t*. This noised data is our input to the neural network.

Since the noise scheduler defines the variance at all timepoints, we also use it to describe the noised data at *t* − 1 from *x*_0_. Recall the neural network is predicting the mean of the reverse kernel. Therefore, given the variance for *t* − 1 predicted by the neural network, we can generate a sample from a constructed distribution and calculate the probability of that sample coming from the true distribution derived from *x*_*t*-1_. This is what the loss represents in Equation 17. During generation, the variance predicted from the model is always used to define the *t* − 1 sample. That sample is then treated as the mean for the next step, but the true noise scheduler variance is used as the next input into the neural network

The Euclidean noise scheduler, *ε*_Euc_, is based on a heuristic 6^th^ order polynomial found to work best in tandem with the other two data distributions. Both the SO(3) and SO(2) schedulers are based on a cosine scheduler proposed by Nichol and Dhariwal(*67*):

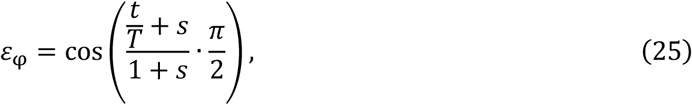

where *s* = 0.008 is an offset used to prevent *ε*_SO(3)_ or *ε*_SO(2)_ from becoming too small near *t* = 0. In the case of *ε*_SO(2)_ we reverse the scheduler owing to the inverse nature of *ρ* (Equation 24).

This segregated approach to applying the various noise schedulers ensures that the different data distributions are treated equally during data destruction and therefore treated as equally vital by the neural network.

### Contact map generation and uses

Contact maps are generated from a noised protein *x_t_*, ***not*** from a noising of the contact maps extracted from *x*_0_. Given the noising of the individual data distributions as described above, we reconstruct an all-atom version of the protein, ignoring hydrogens, and calculate the contact map from that. Specifically, we consider a contact to be defined provided there are any two atoms within 5 Å of one another between two residues. Since the centres of masses of the respective sidechains are then taken to calculate the contact distance, the actual distances used to construct the contact map may be greater than 5 Å, up to a maximum of ∼12 Å for the bigger sidechains at low noise points. Any non-contacts between two residues are set at a maximum distance value of 15 Å, and the contact maps ultimately normalised between 0 and 1 before being input into the neural network. The contact maps are technically redundant with respect to the noised protein input; however, their use aids the network in learning key biochemical relationships, and without their input the structure-sequence relationship is poorly recapitulated. We never predict contact maps with neural network *θ* – they always used purely as ancillary information.

### Network architecture summary

The neural network was designed *ad hoc* to treat each input data type (see Protein Mapping above) as independent streams to emphasise the importance of each designable component, before the hidden dimensions of all are concatenated together and further operations are performed. The penultimate hidden layer is used to project back to all independent data types. We summarise the overall architecture below.

1. The translations, rotations, amino acid vectorisation, backbone and sidechain dihedrals, chain identities, the variances of all data distributions, the current timestep, and the contact maps are all input into the forward pass of the neural network.
2. A time embedding is created via a sinusoidal embedding, followed by a linear projection into a hidden dimension, a GELU activation function, and another linear projection back into the respective feature dimension.
3. Class labels (bound or unbound training samples) are similarly embedded via a simple linear layer and summed to the timestep embedding.
4. Eight hidden states for each data type: coordinate, so(3) vector, amino acid, chain identity, backbone dihedral, sidechain dihedral, and two contact maps are generated by concatenating the input feature, the associated variance (except for in the case of the contact maps), and the timestep embedding.
5. A multilayer perceptron (MLP) is applied to each hidden feature, consisting of a three linear + ReLU activation layers followed by a final linear layer into a larger hidden dimension.
6. A LayerNorm is applied to each hidden feature.
7. A positional encoding based on the sequence length is generated and concatenated to each feature. While the timestep embedding is agnostic of sequence and is applied equally to all features within a sample, the positional encoding may introduce noise if added prior to the MLP layers above. While additional epochs of training would eventually resolve this issue, its inclusion is only vital when attention is applied, hence only including the embedding here.
8. Self-attention is applied to each feature independently. For the rotation and translation invariant inputs, we use the invariant point attention (IPA) introduced by AF2 to ensure invariancy in our model. We use 8 attention heads each with a dimension of 16, applying the transformer successively in four individual instances, each with their own weights and bias. The timestep of the diffusion process (used by the AdaLayerNorm in the IPA), translations and rotations are all included.
9. All output individual features are concatenated into one much larger feature vector with a large hidden dimension describing the whole protein.
10. A positional embedding is again concatenated onto the new hidden feature.
11. Multi-head attention is applied, again with the timestep, translations, and rotations, with the same dimensionality as the self-attention described above.
12. The output is split into six separate streams – all protein mapping data except the contact maps, and another MLP applied similar to that described above.
13. A linear layer is used to project the output from the MLPs back into the dimensionality expected from our data types, for example 3 in the coordinate case or 20 in the amino acid. This output is our final prediction of the *t* − 1 state from our neural network.
14. The predicted variances are extracted by applying a linear layer to the full hidden feature output from the previous MLP timestep. A linear layer is applied for each data distribution, i.e. one for Euclidean, one for so(3), and one for SO(2). We multiply the hidden feature vector by three in the Euclidean case, and by two in the SO(2) case, both based on the number of features described, as we found that predicted superior variances.
15. Finally, a softmax is applied to the three variance output vectors.

We train with a batch size of 64 distributed across two NVIDIA 3090Ti for 400 epochs representing convergence. We apply the AdamW optimizer(*68*) with an initial learning rate of 0.0001, an initial weight decay of 0.01, and a *β*_1_ = 0.9 and *β*_1_ = 0.999. The Exponential Moving Average of weights is used throughout training, and a cosine annealing scheduler is used to adjust the learning rate.

### Conditional constraints

There are multiple conditions that can be included during generation. All conditions can be mixed and matched as needed, and indeed this was the case in the GHR design campaign.

#### Sequence/structure condition

We can design sequences and/or structures by fixing certain features based on some known condition, like a PDB backbone, individual residues, or even chain identity. Since the model is trained to maximise − log *p*(*x_t_*_-1_ |*x_t_*), if all or part of *x_t_* is derived from an input condition, for example some backbone or a partial sequence that needs to not be designed, *x*_t-1_ is predicted within the context of that input condition. Practically, during reverse diffusion we discard the model’s output for any flagged non-designable features and take instead as input for the next timestep the noised version of the clean input conditioning data.

The sharing of logits is used to design homodimers. Specifically, the design pipeline will average the logits across the sequence space for *N* residues, therefore enforcing the sequence to be mirrored across the complex. Crucially, unless requested via backbone fixing, this does not imply any symmetry in the dihedral space, therefore any necessary kinks in the backbone or asymmetric arrangement can be accommodated for.

#### Switch design condition

The sharing of logits can be taken further to design switches. In switch design, we run two or more instances of the diffusion process, and at each timestep share the logits across pre-specified residues or backbone segments. In the GHR example, since the binding interface effectively rotates 180° between the putative inactive and active states, we selected all residues at the interface as designable in each of the respective states. We elected to not simply replicate the feature vector corresponding to the interface from state A to the equivalent non-interface in state B as the implicit membrane should also be accounted for. In other words, one should not expect residues hyperoptimised for a bound configuration to be compatible with a state where they face away from the interface. Instead, to strike a balance between the energetics of binding and unbinding we applied a weighting bias. At each diffusion step the model’s output for the amino acid feature vector in state A is biased towards state B’s output by 20%, and vice versa. Sequence convergence is always achieved at *t* ≈ 460. The bias is applied such that the sum of the vector for any one residue position is still unity. The 80%/20% split was heuristically identified through the post-processing pipeline (see below). It offers the best success rate in terms of oracle structure prediction for two states, enabling switchable behaviour in our designs and avoiding the trapping in either an active or inactive bound state configuration.

### Post-processing validation

We apply a multitude of *in silico* validation techniques as part of our design pipeline. Crucially, this process requires no human intervention yet ensures a high degree of success in designs. In general, we design ∼30000 putative PDBs for each design case of GpA, EphA2, FGFR3, and GHR (× 2 for the two switch states of GHR), and narrow this down to the final 15-30 with the following pipeline. Within the GitHub repo for TMDF we include a detailed README for the pipeline and include easy-to-use scripts to implement each step.

There are four main stages to generating the final designed PDBs:

1. Cleaning the RAW output
2. Validation with either ColabFold(*38*) or ESMfold(*69*)
3. Rosetta relaxation on output structures
4. Calculation of energies for the complex and interfacial potential

Stage 1 removes any output PDBs featuring redundancy in sequence, cysteine or proline, and antiparallel helices (since all designs are for single pass dimeric receptors). It can also check that any desired residue is or isn’t at the interface. This was used in the FGFR3 binder design case to ensure the targeted glutamate was present at the binding site. These different checks can be switched on and off as needed.

In stage 2 we apply ColabFold (ESMfold is also compatible) to mass predict the individual structures using the default settings of 3 seeds and 5 structures per seed, giving 15 conformations per sequence. Flanking regions of the WT sequences are attached to fasta files generated in a pre-processing step, typically we include the flanking region up to the juxtamembrane linker according to a consensus between UniProt(*70*) and DeepTMHMM(*71*). The use of a pre-built MSA in the case of ColabFold and templates are possible as options. In the case of the GHR scaffolds, we parsed a single-sequence MSA and used templates of either the active or inactive states to ensure the correct folding state given the sequences are identical.

Outputs are only accepted if they meet a strict backbone RMSD criteria to the original input PDB used to condition the model. To maintain an ensemble of states and thus rough measure of conformational dynamics, all conformations output by the structure prediction oracle under the RMSD threshold are retained. In the case of GpA and GHR, this threshold was 3 Å to facilitate a diversity of states without straying too far from the native conformation. For GHR, both the active and inactive states needed to be matched for the sequence to be accepted for the first set of 17 final sequences, and only the inactive state in the final set of 13. With EphA2 a more stringent criterion of 1.5 Å was applied owing to the more challenging left-handed nature of the design, and thus lower confidence in the output. Finally, for FGFR3, no RMSD check was needed as it is an inhibitor target.

Owing to the small size of the transmembrane dimers, the small quantity of membrane training data to learn from, and occasionally the *de novo* nature of our designs, we found the typical pLDDT, PAE, and ipTM scores to be poor measures of success. In the case of right-handed helices, they were consistently excellent, even for poor designs or the profiled case of GpA as a trimer or tetramer. The opposite is true in the left-handed case, with all indicating poor structures despite the evidence from both the literature and indeed *a posteriori* from our own TOXGREEN data. Therefore, we elected to apply only the more traditional measures of protein design success via the RMSD and forcefield energy calculations.

Stage 3 involves Rosetta relaxing(*72, 73*) both the bound dimeric and unbound monomeric states of the accepted PDBs. After generating spanfiles, again using a consensus from UniProt and DeepTMHMM, we relax all structures via RosettaScripts(*74*) using the high resolution RosettaMembrane pre-talaris forcefield(*8*), which emphasises critical weak CA-H – O hbonds and bifurcated hbonds without the later introduced bug of hydroxyl hydrogens pointing towards one another leading to an incorrect double satisfaction of a polar group. The equivalent relaxation process is also applied to the initial input PDB used for generation.

In stage 4 we compare the complex and interfacial energies of designs versus the initial input into TMDF. Designs are accepted if: 1) the interfacial energy is better than input, 2) the complex energy is better than input, 3) the sequence identity is lower than a certain threshold, we used 70% for GpA, EphA2, and GHR, although most sequences are below 50% identity, with some as low as 0%. The interfacial sequence identity is also checked to ensure it isn’t above 80%.

A buffer in the complex energy is possible to apply, up to +10REU from the initial input, a value identified by examining the relationship between the complex energy of known GpA mutants and their binding kinetics (see Calculation of Rosetta energies for Binding Kinetics Section in supplemental information). However, in practice so many designs pass the listed criteria that it is more practical to set the buffer to some negative number to ensure the highest quality designs, we used –10 REU for GpA, –5 for EphA2 and 0 REU for GHR. In the GHR case, these criteria were compared with both the unbound and bound states according to the design states respectively. For FGFR3, and in general any pure *de novo* or inhibitor design problem, the top 1% of scoring results in terms of Rosetta energy are returned.

Through this pipeline we identified the final 16, 15, 15, and 30 designs for GpA, EphA2, FGFR3 and GHR respectively. The second round of 13 designs for GHR were extracted by ignoring the success rate of the putative active-like state designs and applying the pipeline only to the inactive state.

### Rosetta design of FGFR3 mutant

The NMR structure of the TM domain of FGFR3 (PDB: 2LZL) provided the starting point for design. The monomeric subunit was extracted and the disordered regions removed, leaving a slice from E370 to R401. The Rosetta CoupledMoves(*75*) protocol was next applied to mutate the A391 position to glutamate, corresponding to the A391E mutation associated with Crouzon syndrome(*49*). CoupledMoves was used with the backrub mover, with a kT of 0.6, neighbourhood repacking, rotamers generated and selected using bump check and energy bias, and a non-fixed backbone over 2000 Monte Carlo cycles. From the 10 trajectories produced, the lowest energy monomer was selected and taken forward for relaxation in an explicit lipid membrane using MD.

### Molecular Dynamics Simulations (GHR)

Starting structures of the GHR designs in both inactive and active states were obtained from the highest scoring Rosetta relaxed structure from the ColabFold pool of predictions, giving 38 initial structures, including the WT. The dimeric complexes were inserted into a 55 nm x 55 nm POPC:POPE:POPG lipid bilayer in a 3:1:1 ratio lipid bilayer and solvated by a 25 nm layer of water above and below the bilayer with 0.15 M of Na^+^ and Cl^−^ ions using the CHARMM-GUI bilayer builder(*76, 77*). Simulations were performed with GROMACS 2024(*78*) with the Colvars module installed(*79*), using the CHARMM36 forcefield(*80*).

For each system we first energy minimized using a steepest descent algorithm until the maximum force converged to < 1000 kJ mol^-1^ nm^-1^, before performing seven rounds of equilibration with decreasing restraints at 310 K, outlines as follows:

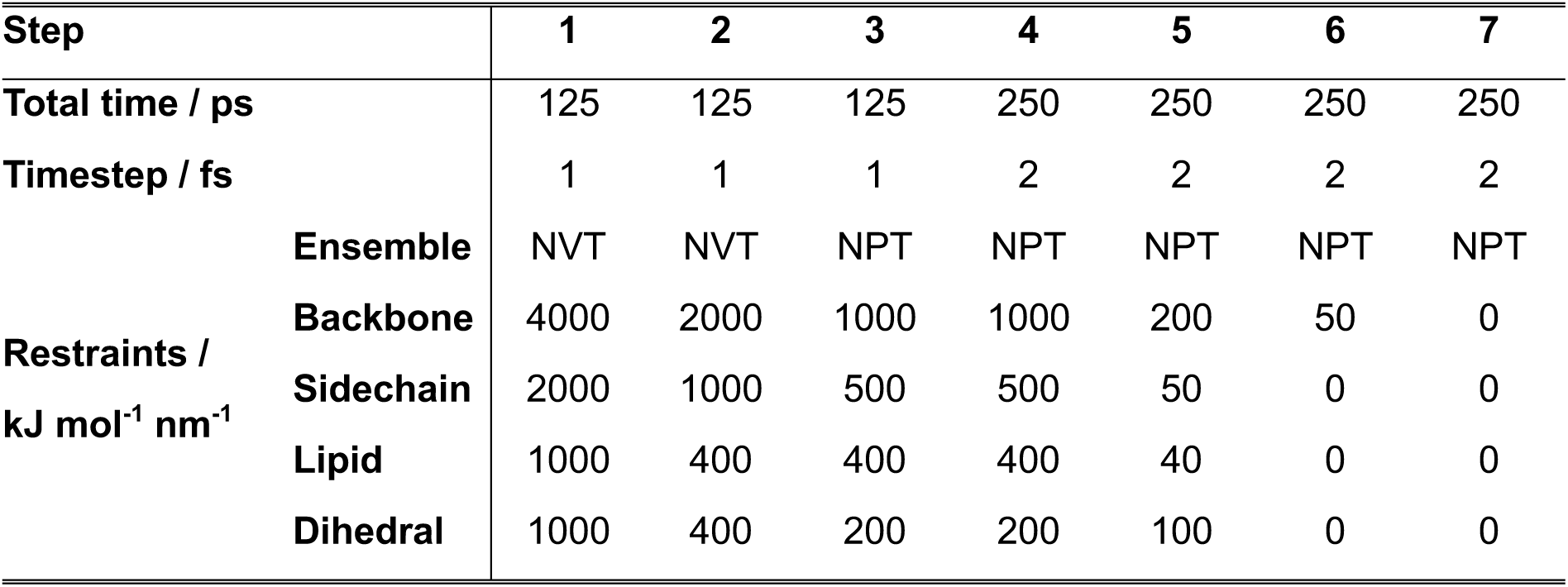

All steps featured a Velocity-rescaled thermostat(*81*) with a timestep of 0.1 ps, with the solute, membrane and solvent coupled independently. The stochastic cell rescaling barostat(*82*) was used for the NPT steps, featuring a semiisotropic pressure coupling time constant of 5.0 ps, a compressibility of 4.5 E-5 bar^-1^, and a reference pressure of 1 bar. A 1.2 nm cutoff distance was selected to consider non-bonded interactions, with PME used for long range interactions.

After equilibration, 1 μs of production time was collected for each system, with the first 50 ns of each ultimately discarded to facilitate equilibration of the backbone. Besides the lack of restraints, production used the same settings as the NPT ensemble steps above.

Following completion of the production runs, a trajectory of each complex was extracted and aligned along the backbone. In the systems where the dimer separated over the course of the simulations (**Figure S2)** frames in the unbound state were removed from the trajectory. A combination of PCA decomposition and KMeans clustering was then performed on all trajectories to discriminate possible conformations of the dimeric state.

### Molecular Dynamics Simulations (FGFR3)

The initial configuration of the monomer taken from the Rosetta mutated form of the NMR structure (PDB: 2LZL), was relaxed through MD using the same settings as above, with a production time of 200 ns. The final frame of the simulation was used as input to TMDF.

### Clustering of Molecular Dynamics simulations (GHR)

For each trajectory, a distogram was initially extracted per frame from the trajectories of the aligned simulation, with each frame representing 1 ns. Distances were calculated between each side chain’s centre of mass (CA in glycine’s case), with the operation only performed between chain A and chain B, i.e. no intrachain contacts were considered. All distances below 8 Å were recorded, with every distance above that capped at a flat 8 Å. Thus, for every design we obtained a distogram with dimension (*T*, *n*_res_, *n*_res_), where *T* is the number of frames, and *n*_res_ the number of residues.

For decomposition by Principal Components Analysis (PCA), each distogram was flattened into a high dimensional vector with shape (*T*, *n*_res_ × *n*_res_) and projected into a lower dimensional space of (*T*, 2) using the Halko et al. method(*83*) implemented in scikit-learn. KMeans clustering was performed directly on the transformed data, with *K* determined by maximising the Calinski and Harabasz score(*84*) between *K* = 2 and *K* = 10, with the objective of having a small internal cluster variance and large between cluster variance. The centres of these clusters are taken as representative conformations from the simulation (**Figure S2**).

### Enhanced Molecular Dynamics Simulations (GHR)

Representative conformations from clustering were taken as input for the steered MD simulations. The combination of GHR1-17 sequences (in addition to the WT) with both active and inactive state GHR, coupled with the number of conformations extracted from the previous step, gave a total of 84 starting structures for the MD.

System setup followed the same procedure as described above for the free dynamics of the GHR designs, and the systems were minimised and equilibrated following the same protocol. The collective variable (CV) for the steered MD production cycle was the inter-helical distance, defined as the distance between the respective centres of mass calculated from the CA of each residue. This distance was projected into the plane of the bilayer, i.e. there was no vectoral component orthogonal to the membrane. The TMs were pulled apart along the CV using the metadynamics method, with the deposited hill weight set to 0.005 kJ mol^−1^, a Gaussian width 2*σ* hill of 2.0 grid points – with a grid point width of 0.2 Å, and hills deposited to the potential every 1000 timesteps. Runs were continued until rupture of the bound state, in the most extreme case of GHR4 this equated to a CV of 3.5 nm, well beyond the equilibrium distance of 0.7 nm, at a free energy of 110 kJ mol^−1^. Given the multiple starting conformations extracted from the free MD, free energies reported in Table S2 are calculated from the average of all conformation’s potential of mean force curves (**Figure 6e**).

## Experimental Methods

### TOXGREEN dimerization assay

Genes encoding the GpA, GHR, and EphA2-based transmembrane domains of interest (Gene Universal Inc.) were digested with NheI and BamHI and ligated into the compatible restriction sites of the pccGFPKAN plasmids (kind gift from Prof. Alessandro Senes, University of Wisconsin-Madison)(*40*). The plasmids were transformed into the MalE-deficient *E. coli* strain MM39 and selected on lysogeny broth (LB) agar plates containing 0.1 mg/mL ampicillin. Three colonies were picked from each construct and grown in selective M9 medium for 18 h at 37 °C. Overnight cultures were added directly to a black 96-well, clear-bottom plate. Absorption at 580/10 nm and GFP fluorescence emission spectra (excitation maximum 485/20 nm and emission maximum at 530/30 nm) were collected using an Infinite 200 PRO plate reader (Tecan). The results are reported as the ratio of fluorescence emission at 530 nm to absorbance at 580 nm, normalized to the wild-type construct.

### DN-AraTM dimerization assay

Genes encoding the FGFR3-based transmembrane domains of interest (Gene Universal Inc.) were digested with SacI and KpnI and ligated into the compatible restriction sites of the pAraTMwt and pAraTMDN (kind gifts from Prof. Bryan Berger, University of Virginia)(*85*). The plasmids and the reporter plasmid (pAraGFPCDF) were co-transformed into the AraC-deficient *E. coli* strain SB1676 (E. coli Genetic Resource Center) and streaked onto selective LB agar plates containing 0.1 mg/mL ampicillin/kanamycin/spectinomycin. Colonies were picked from each construct and grown in LB medium containing selective antibiotics for 6 h at 32 °C. Each culture was diluted in autoinduction medium containing selective antibiotics and grown for an additional 18 h at 30 °C. A series of 2-fold dilutions of the cultures was prepared in a black 96-well, clear-bottom plate. Absorption at 580/10 nm and GFP fluorescence emission spectra (excitation maximum 485/20 nm and emission maximum at 530/30 nm) were collected using an Infinite 200 PRO plate reader (Tecan). The results are reported as the ratio of fluorescence emission at 530 nm to absorbance at 580 nm and normalized to the wild-type heterodimer.

### Maltose complementation test

pAraTMwt and pAraTMDN-based plasmid constructs were transformed into the MalE-deficient *E. coli* strain MM39 and streaked onto selective LB plates 0.1 mg/mL kanamycin. The following day, individual colonies were picked and grown in selective LB medium. Saturated culture from each construct was streaked onto selective M9 minimal medium plates containing 0.4% (w/v) maltose and incubated for 3 days at 37 °C.

### Immunoblot Analysis

Samples were diluted to the same OD_600_, then boiled for 10 min at 95 °C and resolved by SDS-PAGE on a 10% Tris-glycine gel. The samples were transferred to a 0.45-μm nitrocellulose membrane (Cytiva, catalog no. 10600003) using a Trans-Blot Turbo Transfer System (Biorad, catalog no. 1704150). Membranes were blocked with 5% BSA in TBS, Tween 20 (TBS-T) for 1 h at RT. After blocking, the membranes were incubated in anti-maltose binding protein (MBP) at a 1:10,000 dilution (New England Biolabs, catalog no. E8032S) for TOXGREEN constructs and two separate antibodies, anti-HA and anti-Myc, at 1:4,000 and 1:1,000 dilutions, respectively (Invitrogen catalog no. BSM-33157M and MA1-21316) for DN-AraTM at 4 °C overnight. Membranes were then incubated in an anti-mouse HRP-linked antibody (Cell Signaling Technology, catalog no. 7076) at a 1:4,000 dilution for 1 h at RT. Subsequently, membranes were washed with TBS-T. The immunoblot was visualized by chemiluminescence after incubation in Clarity Western ECL substrate (Biorad) using the ChemiDoc XRS+ Molecular Imager (Biorad).

### Construction and functional validation of designed GHR receptors in HEK293T cells

#### Construct design

The native signal peptide of full-length human growth hormone receptor (UniProt ID P10912; residues 1-18 as identified using the SignalP 6.0 webserver(*86*)) was replaced with a murine IgK signal peptide followed by a 3x hemagglutinin (HA) tag ang a GGSGGS spacer. The resulting IgK-3xHA-GGSGGS-GHR(19–638) was inserted into a pcDNA3.1(+) vector. Residues 271-283 (IIFGIFGLTVMLF) of the GHR wildtype transmembrane domain were replaced with the ones designed with TMDF. Plasmids were ordered from GenScript in HT transfection grade quality.

#### Cell culture and transfection

HEK293T wild-type cells were maintained in DMEM (Gibco, #41965-039) supplemented with 10% FBS (Gibco, #10270-106) at 37 °C, 5% CO₂, 95% relative humidity. For ELISA and pSTAT5 assays, tissue-culture–treated 96-well white, clear-bottom plates were used (Corning, #3610); where indicated, a white vinyl backing was applied to the plate underside (Thermo Scientific, #236272).

50’000 HEK293T cells/well were transiently co-transfected with 200 ng pSTAT5 inducible luciferase reporter pGL4.52 (Promega, #E465A) and 5 ng of either GHR wildtype, GHR TMD variants, or empty vector (pcDNA3.1(+)). Cells were first seeded in 100 µL DMEM + 10% FBS in tissue-culture–treated white clear-bottom 96-well plates (Corning, #3610) coated with poly-D-lysine (0.1 mg/mL in ddH_2_O; Sigma, #P6407-5MG). After 1h incubation, 50 µL of the transfection mixture containing 0.5 µL Lipofectamine 2000 (Invitrogen, #11668-019), DNA and Opti-MEM I + GlutaMax-I (Gibco, #51985-026) was added. After 24h post transfection, the medium was exchanged for 150 µL of starvation medium (DMEM without FBS). Following 18-20h of starvation, plates were either subjected to ELISA to measure receptor surface expression, or further used to measure ligand-induced pSTAT5 signaling.

#### Enzyme-linked immunosorbent assay (ELISA) for receptor expression

Cells were fixed with 4% paraformaldehyde in 1x DPBS (Gibco, #14190) (dilution prepared from 16% PFA (EMS, #30525-89-4)) for 10 min at RT and blocked with 2% w/v BSA (Sigma, #A7906) in 1x DPBS for 45 min. Following this, cells were incubated for 30 min each first with mouse anti-HA tag monoclonal antibody (Invitrogen, #26183-1MG) diluted 1:1200 in 2% BSA and then with anti-mouse IgG HRP-linked antibody (Cell Signalling Technology, #7076) diluted 1:500 in 2% BSA. Next, the cells were incubated for 10 min with a 1:1 mix of reagents A and B from the SuperSignal West Pico PLUS kit (Thermofisher, #34577). Following this, the bottom of the plate was covered with white tape (Nunc sealing tape white vinyl, Thermo Scientific, #236272) and luminescence was recorded at the FlexStation3.

#### Measuring growth-hormone induced pSTAT5 signalling & data analysis

Following starvation, the medium was exchanged for 50 µL starvation medium supplemented with or without human growth hormone (PeproTech, #AF-100-40-250ug; Reconstituted to 100 µg/mL aliquots in 1x DPBS, 0.1% BSA). After 4.5h stimulation, of which the last 15 min were at room temperature, 25 µL OneGlo Luciferase buffer (Promega, #E6110) was added. The bottom of the plates was covered with tape and after 5 min of incubation at room temperature luminescence was measured using the FlexStation3.

For basal versus ligand-induced signaling, cells were stimulated with 250 ng/mL GH or vehicle (0 ng/mL). For dose–response analysis, twofold serial dilutions of GH were prepared from 500 ng/mL to 7.8 ng/mL, including a 0 ng/mL condition. All assays were performed in three independent biological replicates, each measured in technical triplicate. Within each experiment, signals were averaged across technical replicates, background-subtracted using empty vector controls (only for basal versus ligand-induced signaling), and normalized to the stimulated wild-type control (250 ng/mL for basal vs stimulated assays; 500 ng/mL for dose– response curves). Normalized data from independent experiments were pooled. Nonlinear regression was applied in GraphPad Prism 10.2.0 (Dose–response – Stimulation, *[Agonist] vs. response -- Variable slope (four parameters)* model, default settings).

### Designed TM peptide production for X-ray crystallography of GpA

Peptides were expressed recombinantly in *E. coli* as 9His–trpLE fusion proteins following a modified version of the procedure described by Sharma *et al*(*87*). Briefly, fusion proteins were solubilized from inclusion bodies, purified using HIS-Select Nickel Affinity Gel (Merck), subjected to cyanogen bromide cleavage and peptides were purified by reverse-phase HPLC using a ZORBAX StableBond 300 C3 column (Agilent). For cysteine-containing peptides, the C-terminal cysteine sulfhydryl group was protected during cell lysis and inclusion body solubilization by the addition of 10 mM S-methyl methanethiosulfonate (MMTS, Sigma-Aldrich). At no stage were peptides allowed to form disulfide bonds. Final HPLC-purified products were lyophilized and stored at room temperature until required.

### SDS-PAGE analysis for X-ray crystallography of GpA

Gel samples were prepared by taking indicated amounts of each purified peptide (**Figure Sb**) from dried and weighed product redissolved in 1,1,1,1,1,1-hexafluoroisopropanol (HFIP, Merck). Samples were lyophilized, redissolved in 25 μl 1× NuPAGE LDS sample buffer (Thermo Fisher Scientific), and heated for 1 min at 95°C. Cooled samples were separated on 12% NuPAGE Bis-Tris gels (Thermo Fisher Scientific) at 200 V for 35 min and visualized by staining with Coomassie Blue R-250 (Bio-Rad) (**Figure S4a**).

### Crystallization screening and data collection for GpA designs

For reconstitution into LCP, lyophilized peptide was weighed and co-dissolved with appropriate amounts of monoolein (Nu-Chek Prep) in HFIP. Solvent was removed under streaming nitrogen, followed by lyophilization overnight. Peptide-monoolein mix was heated (52°C) until molten and mixed 3:2 with 10 mM Tris pH 8.0 for LCP formation using coupled 100 μl gastight Hamilton syringes (Formulatrix) at room temperature.

For crystallization screening, LCP was dispensed in 100 nl drops onto 96-well glass plates (Molecular Dimensions) with 1000 µl of precipitant solution using a Mosquito LCP robot (TTP Labtech) at room temperature. Plates were sealed and incubated at 20°C in a Rock Imager 1000 (Formulatrix) for automated imaging and monitoring of crystal formation. All screening was performed at the Melbourne Protein Characterisation facility (Bio21 Institute, University of Melbourne) using JBScreen LCP HTS (Jena BioScience), MemMeso (Molecular Dimensions), and PACT Premier (Molecular Dimensions) screens.

A version of GpA3 was produced without the N-terminal Glu-Pro-Glu and the C-terminal cysteine to improve on an initial crystal hit that diffracted poorly. This peptide crystallised in 0.1 M MES pH 6.0, 0.01 M copper (II) chloride, 0.2 M ammonium formate, and 25% (v/v) PEG DME 500 and was used for structure determination. Diffraction data were collected on the MX2 beamline of the Australian Synchrotron(*88*) at a wavelength of 0.9537 Å and a temperature of 100 K.

### Data analysis and structure determination for GpA3

Diffraction images were indexed and integrated with DIALS. Two independent lattices were detected in the dataset. The second lattice was related to the first by the change-of-basis operator:

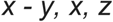

Rotation matrix to transform crystal 1 to crystal 2:

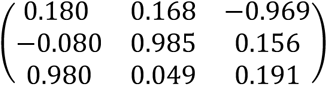

This corresponds to a rotation of −79.742 degrees about the axis:0.054 0.991 0.126 Spotfinding and integration were performed using the max_lattice=2 option in DIALS. Statistics for crystal_0 were better than crystal_1. Frames 160-2000 (0.1° per frame) from crystal_0 were selected as a wedge for scaling. POINTLESS and DIALS symmetry analysis consistently indicated P6222. Merging and outlier rejection were performed with AIMLESS and ctruncate. The merged dataset had strong diffraction to 1.80 Å with I/sigma(I) in the outershell at 3.84 (99.61 percent complete; multiplicity 6.8; overall I/sigma(I) = 17.65; CC1/2 = 1.000; Rmerge = 0.04406). The highest-resolution shells showed the expected fall-off in I/sigma(I). Despite being collected as a native dataset (wavelength: 0.9536), Aimless indicated significant anomalous signal extending to 2.14 Å.

#### Phasing, Model building and refinement (Table S1)

Molecular replacement with Phaser in P6222 with a model helix yielded a unique solution (LLG = 398; TFZ = 17.1). Initial refinement produced interpretable maps, but R-free over 0.44. The data was cut to 2.4 Å for initial rounds of refinement yielding an Rfree of 0.3498. Iterative cycles of manual rebuilding in Coot and restrained refinement in phenix.refine were performed. Strong positive difference density near the N-terminus indicated a bound metal; this was modeled as a single Cu^2+^ ion disordered over two sites separated by 1.55 Å with equal occupancies. The presence and placement of Cu explained the observed anomalous differences. Independent experimental phasing (HySS substructure search followed by Phenix AutoSol SAD phasing in P6222) recovered the same Cu sites and confirmed the MR solution.

Translation–libration–screw (TLS) refinement (per-chain groups) reduced Rfree from 0.3201 to 0.2752 at 2.40 Å. Extending the refinement to 1.80 Å did not further improve cross-validation statistics (Rfree remained 0.29 or higher); the final model was refined against data to 1.8 Å with isotropic B factors, TLS and anomalous groups. All residues were built with continuous density; Cu was refined with two alternative positions and occupancy refined to 0.6/0.4.

#### Pathology assessment and space-group checks

The high R-free in our final model suggests crystal pathology. Potential pathologies were evaluated systematically. Xtriage reported no compelling evidence for merohedral twinning. Re-refinement in lower symmetries (e.g., P6122, P3212, P3221, P32, C2, P1) did not yield substantially improved statistics or maps. Zanuda did not suggest a better lattice or symmetry reduction. Trial refinements with candidate twin laws in P3 space groups produced only marginal decreases in R-free (about 2–3 percent) without concomitant map improvement.

Translational non-crystallographic symmetry (tNCS) was detected along the c-axis (0, 0, 0.291; 24.91%; p-value 3.49e-03) despite there being only a single protein chain in the asymmetric unit. In this crystal form, the c-axis corresponds to the stacking direction of the lamellar layers formed by the lipidic cubic phase (LCP) during crystallization. Within each layer, ordered lipid molecules and associated aqueous channels create alternating hydrophobic and hydrophilic strata. We were able to model one well-ordered monoolein molecule and observed weak, discontinuous density consistent with an additional, partially disordered monoolien. Non-uniformity in the composition and thickness of these lipid and aqueous layers is not well captured by standard bulk solvent models and likely contributes to the elevated R-free values often observed in membrane protein crystals grown in LCP.

## Acknowledgments

We thank members of the Barth lab for helpful discussions.

## Authors contributions

L.S.P.R., and P.B. designed the project; L.S.P.R. and M.S. developed the computational framework and analyzed the results; L.S.P.R. and M.W. designed, selected TM proteins and ran MD simulations; R.B. performed experimental validation of the designed GHR; V.A. performed the TM association experiments under the supervision of D.T. J.V.N. carried out the structural validation of the designs under the supervision of M.J.C. and M.E.C.; L.S. performed experimental validation of the designed TMs; all authors participated in the analysis and interpretation of the results; L.S.P.R., R.B. and P.B. wrote the manuscript with contributions from the other authors; P.B. supervised the entire project.

## Data and materials availability

All data are available in the manuscript or the supplementary materials.

## Supplemental Information

### Supplemental Figures

**Figure S1:**
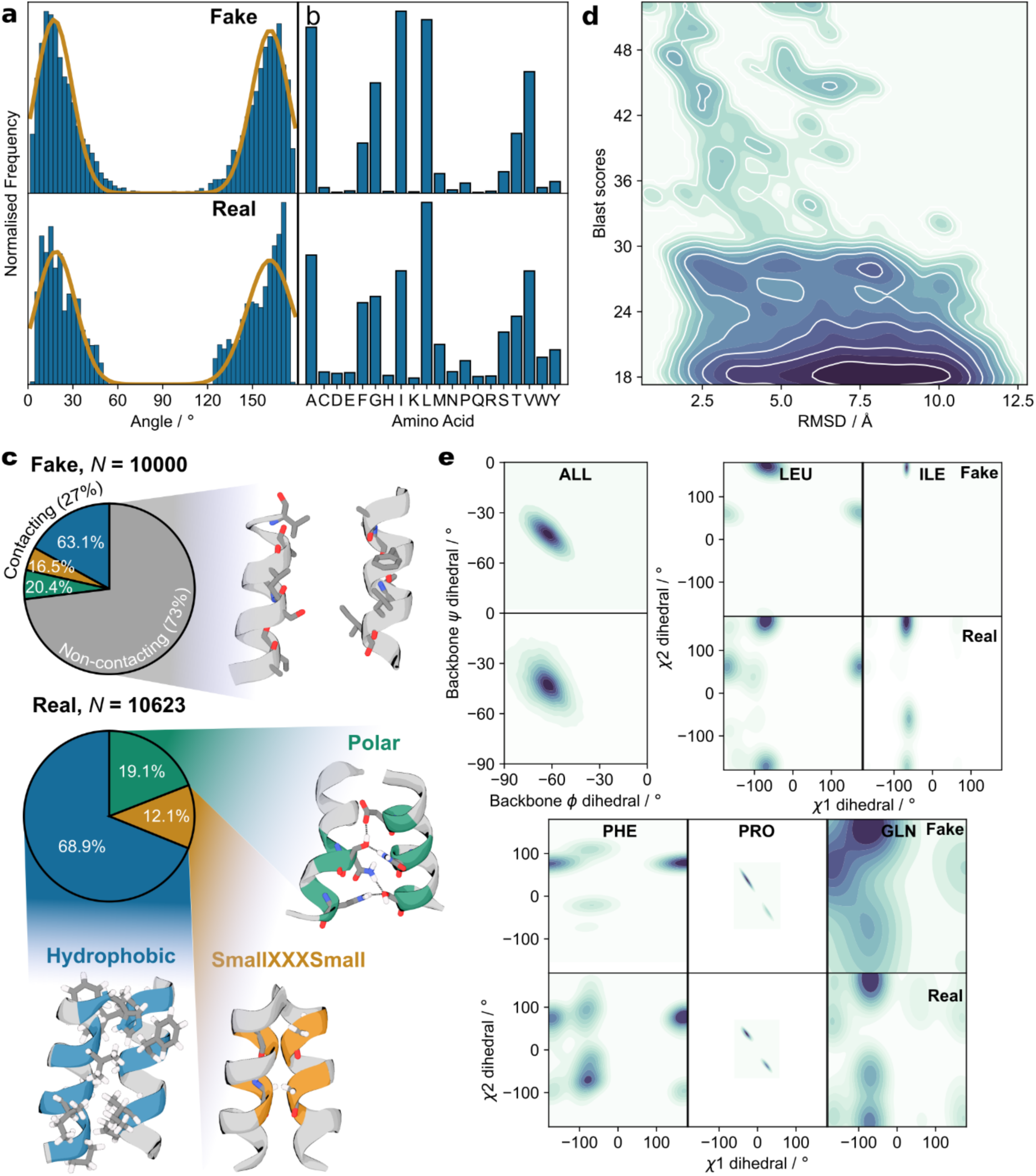
Impact of the ablation of the contact maps for neural network input. (**a**) The crossing angle space between the ablated trained network’s output and the real data is almost identical to the non-ablated data, with the two distributions neatly matching. (**b**) The ablated output feature similar undersampling of polar amino acids to the non-ablated version of TMDF. The key difference here is the significant oversampling of leucines versus that seen in the training data, a problem not mirrored by the non-abated version of TMDF. (**c**) In sharp contrast to non-ablated network, contact motif recapitulation is almost completely non-existent, with most dimers barely featuring even a single designed contact, demonstrating the added value of the contact maps. (**d**) The BLAST vs RMSD relationship of the ablated TMDF is significantly worse than the non-ablated form, demonstrating that not only are contact maps key in describing the interface, but also aid with learning the sequence structure relationship. (**e**) Backbone dihedrals are neatly recapitulated despite ablation of the contact maps, while sidechain dihedral distributions are marginally, though notably, worse to those seen in the real non-ablated version of TMDF.

**Figure S2:**
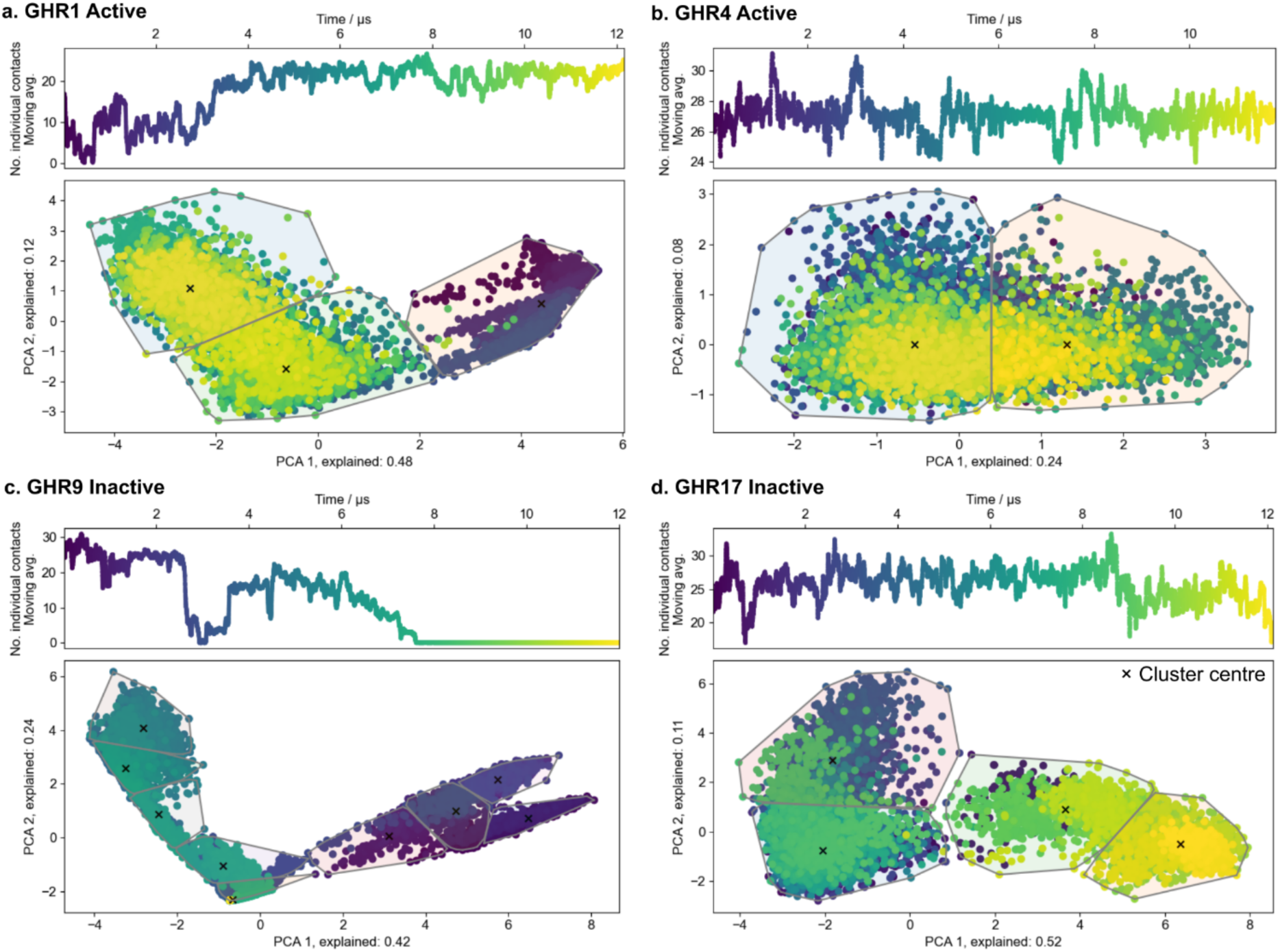
Example summary for the breakdown of the free Molecular Dynamics. Performed for (**a**) GHR1 active (**b**) GHR4 active (**c**) GHR9 inactive (**d**) GHR17 inactive. (**Top**) Count for the number of encountered contacts between chains during the course of the simulation, and (**bottom**) Principle Component Analysis decomposition of the CA coordinates into a 2-dimensional space, from which cluster centres are extracted as the starting conformations for steered MD. Points are coloured by their progress through the simulation in nanoseconds.

**Figure S3:**
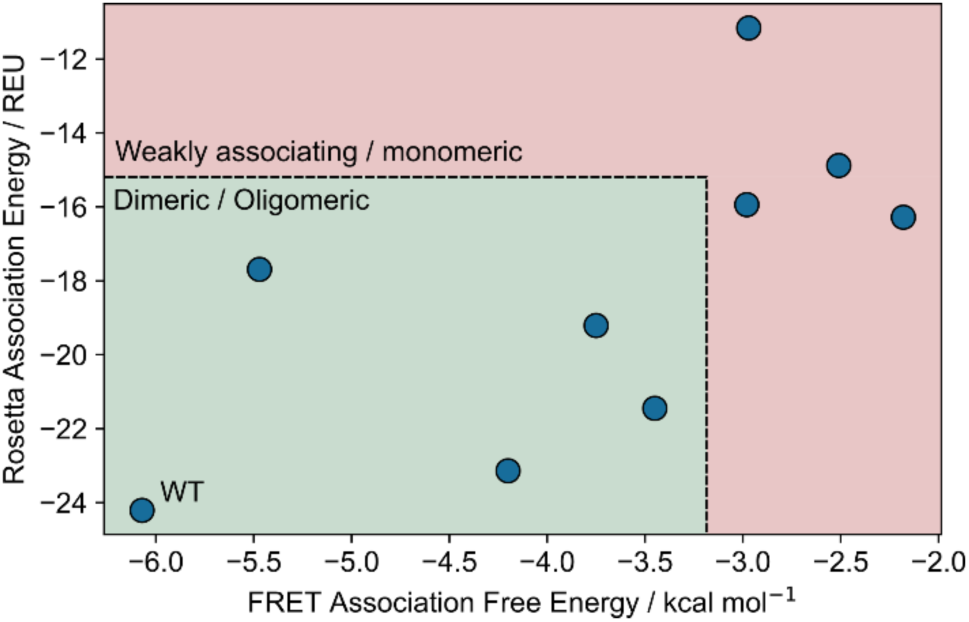
Correlation between the FRET association free energy as measured by Vázquez *et al.*(*37*) versus the Rosetta dimerisation energies for those same mutants. The mutants are split roughly into weakly associating / monomeric and dimeric / oligomeric, forming the threshold (+10 REU vs WT) to consider a TMDiffusion design still in the predicted dimer state. This association also validates our Rosetta guided filtering approach.

**Figure S4:**
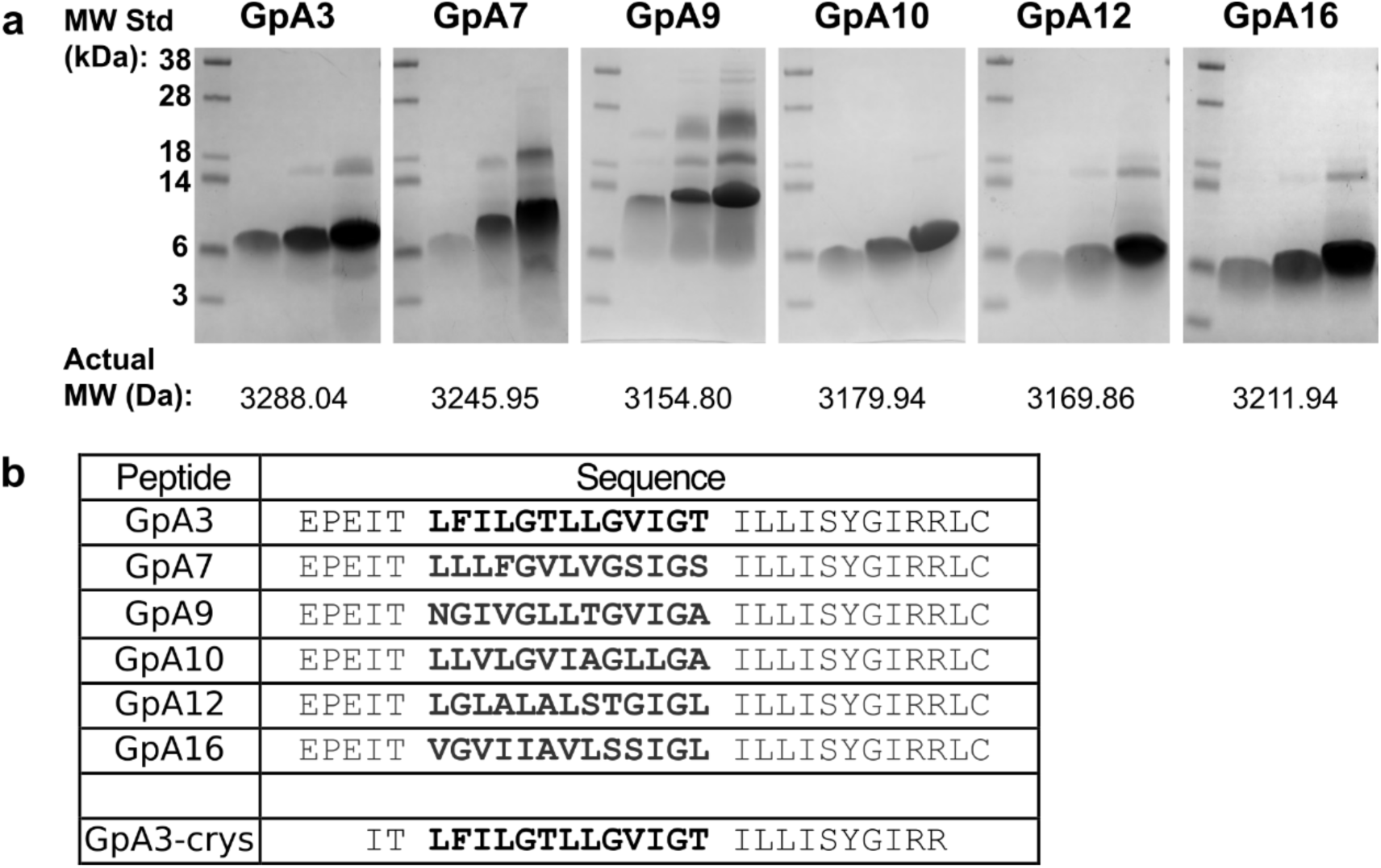
(a) SDS-PAGE. Of GpA designs 3, 7, 9, 10, 12, and 16, with GpA3 corresponding to the resolved crystal structure. 5, 15 and 45 µg of peptide was loaded for each sample. (**b**) Summary of the purified peptide sequences. Bold letters indicate the designed portion of the sequence.

## Supplemental Data

**Table S1.** Data collection and refinement statistics.

|  |  |
| --- | --- |
|  | helix* |
| Wavelength | 0.9536 |
| Resolution range | 27.28 - 1.8 (1.98 - 1.8) |
| Space group | P 62 2 2 |
| Unit cell | 31.505 31.505 117.665 90 90 120 |
| Total reflections | 41365 (8621) |
| Unique reflections | 6069 (1525) |
| Multiplicity | 6.8 (5.7) |
| Completeness (%) | 99.61 (99.28) |
| Mean I/sigma(I) | 17.65 (3.84) |
| Wilson B-factor | 29.08 |
| R-merge | 0.04406 (0.2293) |
| R-meas | 0.04767 (0.2531) |
| R-pim | 0.01783 (0.1032) |
| CC1/2 | 0.999 (0.999) |
| CC* | 1 (1) |
| Reflections used in refinement | 3655 (878) |
| Reflections used for R-free | 367 (88) |
| R-work | 0.2549 (0.2837) |
| R-free | 0.2900 (0.3140) |
| Number of non-hydrogen atoms | 227 |
| macromolecules | 200 |
| ligands | 27 |
| solvent | 0 |
| Protein residues | 26 |
| RMS(bonds) | 0.003 |
| RMS(angles) | 0.60 |
| Ramachandran favored (%) | 100.00 |
| Ramachandran allowed (%) | 0.00 |
| Ramachandran outliers (%) | 0.00 |
| Rotamer outliers (%) | 4.55 |
| Clashscore | 3.99 |
| Average B-factor | 48.05 |
| macromolecules | 47.32 |
| ligands | 53.40 |

**Statistics for the highest-resolution shell are shown in parentheses.**

**Table S2:** GHR Steered MD Barriers. Summary of GHR designs 1-17 steered MD measured energy barriers, where both active and inactive designs were designed for via TMDF’s switch design. Energy barrier is defined as the peak of the potential of mean force. Reported are the averages in peak PMF across all starting conformations. Also provided are the direct comparisons with the corresponding basal activity and EC50 measurements. For reference, body temperature *k_B_T* (i.e. thermal fluctuations), is about 2.6 kJ mol^−1^.

| System | EV | WT | GHR1 | GHR2 | GHR3 | GHR4 | GHR5 | GHR6 | GHR7 | GHR8 | GHR9 |
| --- | --- | --- | --- | --- | --- | --- | --- | --- | --- | --- | --- |
| Basal Activity | 1775 | 2967 | 2400 | 3120 | 4430 | 30200 | 3260 | 2440 | 2610 | 3272 | 3630 |
| EC50 / ng ml<sup>-1</sup> | N/A | 66 | 52 | 69 | 50 | 81 | 65 | 65 | 66 | 65 | 57 |
| Active state barrier / kJ mol<sup>-1</sup> | N/A | 48 | 28 | 110 | 8 | 110 | 7 | 38 | 7 | 48 | 9 |
| Inactive state barrier / kJ mol<sup>-1</sup> | N/A | 9 | 25 | 16 | 30 | 46 | 8 | 14 | 14 | 9 | 9 |

| System | EV | WT | GHR10 | GHR11 | GHR12 | GHR13 | GHR14 | GHR15 | GHR16 | GHR17 |
| --- | --- | --- | --- | --- | --- | --- | --- | --- | --- | --- |
| Basal Activity | 1750 | 3190 | 3460 | 3740 | 3940 | 4350 | 3440 | 9540 | 6030 | 4447 |
| EC50 / ng ml<sup>-1</sup> | N/A | 66 | 76 | 55 | 122 | 59 | 60 | 65 | 81 | 83 |
| Active state barrier / kJ mol<sup>-1</sup> | N/A | 48 | 7 | 24 | 35 | 22 | 24 | 83 | 93 | 47 |
| Inactive state barrier / kJ mol<sup>-1</sup> | N/A | 9 | 12 | 10 | 25 | 21 | 77 | 160 | 91 | 68 |

## Ablation studies

We applied two rounds of ablation to examine the strengths of our approach:

1) A version of the network without transformers

2) A version of the network without contact maps

Both versions of the model were trained to Epoch 400 in equivalency to the standard model.

In the first instance without transformers, the trained model failed to produce any meaningful output. While generated proteins are not random noise, they do not form any correct topology, therefore demonstrating the necessity of transformers for learning long range information across the hidden state.

The model without contact maps was able to produce valid looking samples. However, looking further into the statistics of generated output in an equivalent manner to Figure 2 (**Figure S1**), we observe stark differences between the neural networks. While both the crossing angle distribution and amino acid distributions are similar (**Figure S1a/b**), the number of valid contacts extracted from the generated output is roughly five times smaller *versus* the native TMDF for an equivalent number of 10000 samples (**Figure S1c**). The impact of this is observed in the poor sequence structure relationship learnt by the non-contact map version of the model (**Figure S1d**). While high BLAST scores do tend to correspond to smaller RMSDs, the space is sparsely populated when compared with the normal model (**Figure 2d**). Finally, while the dihedral space is recapitulated by the contact mapless model, the learnt distributions are further from the ground truth (**Figure S1e**).

These results indicate that while some critical features; crossing angles, amino acid distributions, backbone structure etc., are learnt by a model trained only implicitly on contacts via our vectorised protein, the explicit inclusion of contacts aids significantly in emphasising the critical interhelical contacts needed to maximise binding and in the task of associating specific sequence motifs with structural equivalents. Given our objective of emphasising biophysical information to achieve optimised binding, it is clear that the inclusion of contact maps is necessary to achieve high quality outputs from the model.

## Calculation of Rosetta energies for binding proxy

To establish a baseline for the selection criteria of designs based on Rosetta energies, we took data from Vázquez et al.(*37*) who calculated the free energy of dimer association of GpA WT and 9 deleterious mutants with FRET and calculated corresponding Rosetta interfacial energies for each (**Figure S3**). Specifically, we first predicted the arrangement of each with AlphaFold2-multimer(*89, 90*) using templates of GpA WT (PDB: 1AFO), in an approach similar to our post-processing validation pipeline (see Methods). To account for the significant increase in leucines now proximal to the interface in many of the mutants, we introduced minor perturbations to the bound state via RosettaDock(*91*), before relaxing both dimeric and monomeric subunits following out pipeline. The final reported energies are extracted from the average of these calculations.

Figure S3 demonstrates a clear relationship between the association energy calculated from FRET and from Rosetta (*R* = 0.65). Given that mutants G83I, 1A31, 1A32-2, and MCL1 are all either weakly associating or predominantly monomeric at Δ*E* ≈ +10 REU with respect to WT, we established +10 REU as our maximum energy difference to accept samples during the post-processing pipeline. In practice, the difference is normalised to sequence length to account for how Rosetta calculates energies.

